# Dynamic *in vivo* mapping of the methylproteome using a chemoenzymatic approach

**DOI:** 10.1101/2022.07.22.501130

**Authors:** Jonathan Farhi, Benjamin Emenike, Richard S. Lee, Christian M. Beusch, Robert B. Jones, Ashish K. Verma, Celina Y. Jones, Maryam Foroozani, Monica Reeves, Kirti Sad, Kiran K. Parwani, Pritha Bagchi, Roger B. Deal, David J. Katz, Anita H. Corbett, David E. Gordon, Monika Raj, Jennifer M. Spangle

## Abstract

Dynamic protein post-translation methylation is essential for cellular function, highlighted by the essential role of methylation in transcriptional regulation and its aberrant dysregulation in diseases including cancer. This underscores the importance of cataloging the cellular methylproteome. However, comprehensive analysis of the methylproteome remains elusive due to limitations in current enrichment and analysis pipelines. Here, we employ an L-Methionine analogue, ProSeMet, that is chemoenzymatically converted to the SAM analogue ProSeAM in cells and *in vivo* to tag proteins with a biorthogonal alkyne that can be directly detected via LC-MS/MS, or functionalized for subsequent selective enrichment and LC-MS/MS identification. Without enrichment, we identify lysine mono-, di-, and trimethylation, histidine methylation, and arginine methylation with site specific resolution on proteins including heat shock protein HSPA8, for which methylation is implicated in human disease. With enrichment, we identify 486 proteins known to be methylated and 221 proteins with novel methylation sites encompassing diverse cellular functions. Systemic ProSeMet delivery in mice pseudomethylates proteins across organ systems with blood-brain barrier penetrance and identifies site-specific pseudomethylation *in vivo* with LC-MS/MS. Leveraging these pipelines to define the cellular methylproteome may have broad applications for understanding the methylproteome in the context of disease.

## Introduction

Protein methylation is a widely occurring post-translational modification (PTM) that plays an important role in cell signaling, development, and disease^1^. Methylation most commonly occurs on the basic amino acids lysine (Lys), arginine (Arg), and histidine (His): lysine residues can be mono-, di-, or tri-methylated, arginine residues can be monomethylated or dimethylated symmetrically or asymmetrically, and histidine is monomethylated at both the 1 and 3 positions of the imidazole ring^1,2^. On proteins, these methylation events are catalyzed by methyltransferase enzymes using the metabolite *S*-adenosyl-L-methionine (SAM)^3^. SAM-dependent methyltransferases fall into three structurally distinct classes: Class I enzymes contain a conserved seven-stranded β-sheet and include all canonical arginine methyltransferases (PRMTs) as well as the lysine methyltransferase DOT1L; Class II enzymes contain a SET domain and encompass all other lysine methyltransferases (KMTs), and Class III enzymes are membrane-associated methyltransferases^4^.

Histone protein methylation affects epigenetic regulation of gene expression by mediating the recruitment of chromatin modifying enzymes to chromatin and transcriptional regulators to gene regulatory regions including promoters and enhancers^2,5^. Non-histone protein methylation is likewise an essential PTM affecting cellular biology and disease pathogenesis^6,7^. For example, methylation of the PI3K effector AKT by SETDB1 increases the duration of AKT activation, promoting tumor growth in murine models^8,9^. Similarly, methylation of CRAF at R100 by PRMT6 interferes with RAS/RAF binding, suggesting a mechanism by which PRMT6 loss increases tumor initiation and metastasis in hepatocellular carcinoma^10^. Moreover, the oncogenic role of Enhancer of Zester Homolog 2 (EZH2) is hypothesized to extend beyond methylation of histone H3 on lysine 27 (H3K37) to include cytosolic targets: the transcription factors STAT3^11^, GATA4 (K299)^12^; and FOXA1 (K295)^13^, among others. The important roles these methyltransferase enzymes play in normal cellular function and disease has prompted the clinical development of several inhibitors, including the recent first-in-class EZH2 inhibitor tazemetostat for the treatment of epithelioid sarcoma and follicular lymphoma^14^ and several PRMT1/5 and CARM1 inhibitors^15,16^.

Current efforts to identify histone and non-histone protein methylation often employ mass spectrometry (MS) or antibody-based approaches to identify methylated residues and changes to their methylation status^6^. However, unlike acetylation or phosphorylation, methylation does not change the charge or physiochemical properties of the amino acid residue, rendering MS-based characterization challenging without isotopic labeling, chemical modification, or biased enrichment to reduce the complexity of analysis^1,17^. The development of pan-methyl, pan-methyl-lysine, and pan-methyl-arginine specific antibodies has underscored the breadth of the methylproteome^18,19^ but has been limited by cross-reactivity (e.g., between methylated Lys and Arg residues)^20^, batch-to-batch irreproducibility, and an inability to identify novel methylation sites^21,22^. Furthermore, MS-based methods aimed at detecting methylation must overcome high false discovery rates because a high number of amino acid combinations produce sequences that are isobaric to methylated peptides of a different sequence (e.g., the mass difference between Asp and Glu is equivalent to a single methylation event)^23^.

The challenges presented by current MS– or antibody-based approaches have prompted the development of chemical strategies to selectively modify methylated residues^24^ or hijack cellular machinery for a tag-and-modify approach^25^. In particular, SAM analogues have been employed to “pseudomethylate” (tag) proteins using engineered methyltransferases in live cells or on cell lysate^26–30^. Recently, treatment of cell lysates with the SAM analogue propargyl-*Se*–adenosyl-L-selenomethionine (ProSeAM) enabled profiling of the methylproteome and of methylhistidine targets of the histidine methyltransferase METTL9^31,32^. While these *in situ* methodologies demonstrate the ability of selected methyltransferases to methylate a substrate based on a consensus sequence, *in situ* studies cannot capture changes to the methylproteome in response to biological stimuli or reflect methyltransferase activation and localization into unique cellular compartments. Likewise, strategies involving engineered enzymes are unable to profile the broad spectrum of the methylproteome and are therefore restricted to specific applications.

These challenges have prompted the development of novel strategies to chemically label methyltransferase targets in live cells. The L-Met analogue propargyl-L-selenohomocysteine (ProSeMet) is chemoenzymatically converted by cellular methionine adenosyltransferase (MAT) enzymes to the SAM analogue ProSeAM *in vitro.* In turn, ProSeAM is used by native RNA methyltransferases to propargylate RNA with a biorthogonal alkyne tag allowing for downstream analysis^33–35^. Previous reports have not, however, explored the ability of ProSeAM to propargylate proteins using endogenous methyltransferases. As such, we envisioned that dynamic profiling of the methylproteome could be accomplished using ProSeMet through the chemoenzymatic approach (**Fig. 1**). In this work, we describe the application of ProSeMet for methylproteome profiling *in vitro* and *in vivo*. Using ProSeMet, we performed an unbiased analysis of the methylproteome with and without enrichment in the SMARCB1-mutant, EZH2-activated, cell line G401^36^. Prior to enrichment, we identified arginine, lysine, and histidine methylation with site specific resolution, including heat shock protein HSPA8 at R469. Following functionalization and enrichment, we identified 486 previously characterized methylated proteins and the discovery of 221 novel methylated proteins. Furthermore, we demonstrate that *in vivo* ProSeMet administration to a variety of living organisms including *A. thaliana, C. elegans, S. cerevisiae, and M. musculus* resulted in protein propargylation. Specifically, ProSeMet administration to BALB c/J mice leads to labeling and identification of site-specific propargylation via LC-MS/MS in all organ systems analyzed, including the heart, lungs, and brain. Our work demonstrates the development of an improved chemoenzymatic platform that has the potential to study dynamic changes to the methylproteome in response to biological stimulus and disease pathogenesis both in cell and murine models.

**Figure 1.**
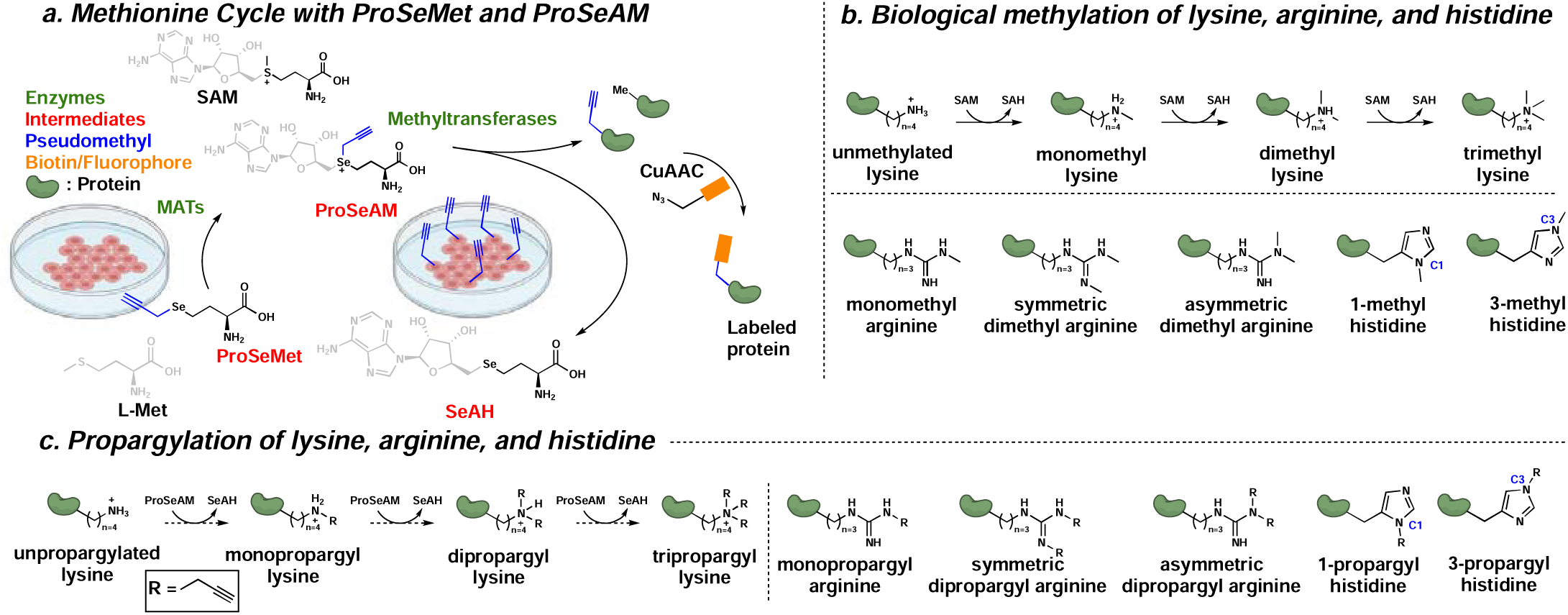
Chemoenzymatic approach for metabolic MTase labeling. **a**. ProSeMet can be converted to ProSeAM by MAT enzymes in live cells. ProSeAM can then be used by diverse methyltransferases to propargylate target protein. **b**. (*top*) Conversion of unmethylated lysine to mono, di, and trimethyl Lys through the action of SAM. (*bottom*) The guanidine moiety on Arg can undergo monomethylation, then symmetric or asymmetric dimethylation. The imidazole ring of His can undergo monomethylation at the C1 or C3 position **c**. (*left*) Conversion of unpropargylated Lys to mono, di, and tripropargyl lysine could occur through the intermediacy of ProSeAM. (*right*) Predicted Arg monopropargylation, symmetric dipropargylation, and asymmetric dipropargylation as well as His propargylation at the C1 and C3 positions.

## Results

### ProSeMet competes with L-Met to propargylate proteins in the cytoplasm and nucleus

Previous literature has reported the use of ProSeMet as a viable substrate of the unaltered RNA methyltransferase METTL3-METTL14 in MOLM13, HEK293T, and HeLa cells^33,35^. We therefore hypothesized that ProSeMet could also be used to deposit a biorthogonal alkyne tag (pseudomethylate) on proteins in living cells using native methyltransferases (**Fig. 2a)**. Protein extracted from T47D cells incubated with ProSeMet subjected to copper-catalyzed azide-alkyne cycloaddition (CuAAC) to attach a fluorescent picolyl azide and separated via SDS-PAGE showed protein labeling that was detectable with as little as 5 µg input protein (**Fig. 2b**). Labeling efficacy scaled with increasing concentrations of ProSeMet until 100 µM concentrations (**Fig. 2c**). Since previously reported L-Met analogues have been shown to incorporate into proteins during translation^37^, we collected lysates from cells treated with L-Met, ProSeMet, or the L-Met analogue azidohomoalanine (AHA) in the presence of the protein synthesis inhibitor cycloheximide (CHX). CHX treatment abrogated AHA labeling but did not affect ProSeMet signal, indicating that the majority of ProSeMet is not incorporated into newly synthesized proteins but rather converted to ProSeAM and used by native methyltransferases (**Fig. 2d**). Furthermore, competition with the unlabeled natural metabolite L-Met revealed a dose-dependent reduction of ProSeMet labeling with almost complete depletion of ProSeMet signal at 10 µM L-Met concentrations (**Fig. 2e**). Since protein methylation occurs in both the cytoplasm and nucleus, we next evaluated the cellular compartmentalization of ProSeMet-directed labeling in live cells. Cell fractionation of the cytosolic and nuclear compartments followed by SDS-PAGE fluorescent analysis revealed no fluorescent labeling of the L-Met control but robust ProSeMet labeling of protein across molecular weights in both cytosolic and nuclear fractions (**Fig. 2f**). We further performed immunofluorescence (IF) microscopy analysis of T47D, LNCaP, and MCF10A cells incubated with ProSeMet or L-Met. Cells treated with 100 µM ProSeMet or L-Met for 16 h were washed to remove unreacted ProSeMet, fixed, permeabilized, and ligated to a fluorophore-conjugated azide. Cells treated with ProSeMet exhibited labeling in both the cytosolic and nuclear compartments, while cells treated with L-Met remained unlabeled (**Fig. 2g, Fig. S1**). Evaluation of condensed chromatin in mitotic cells suggest ProSeMet-directed labeling of DNA is not achieved using this approach (**Fig. S2**). To ensure that ProSeMet would not cause cellular death during the experimental timeframe, we performed Annexin V/PI staining to assess the viability of cells treated with ProSeMet over a time course. ProSeMet treatment did not increase cytotoxicity compared to DMSO treatment at low (50 µM) or high (200 µM) concentrations (**Fig. S3**).

**Figure 2.**
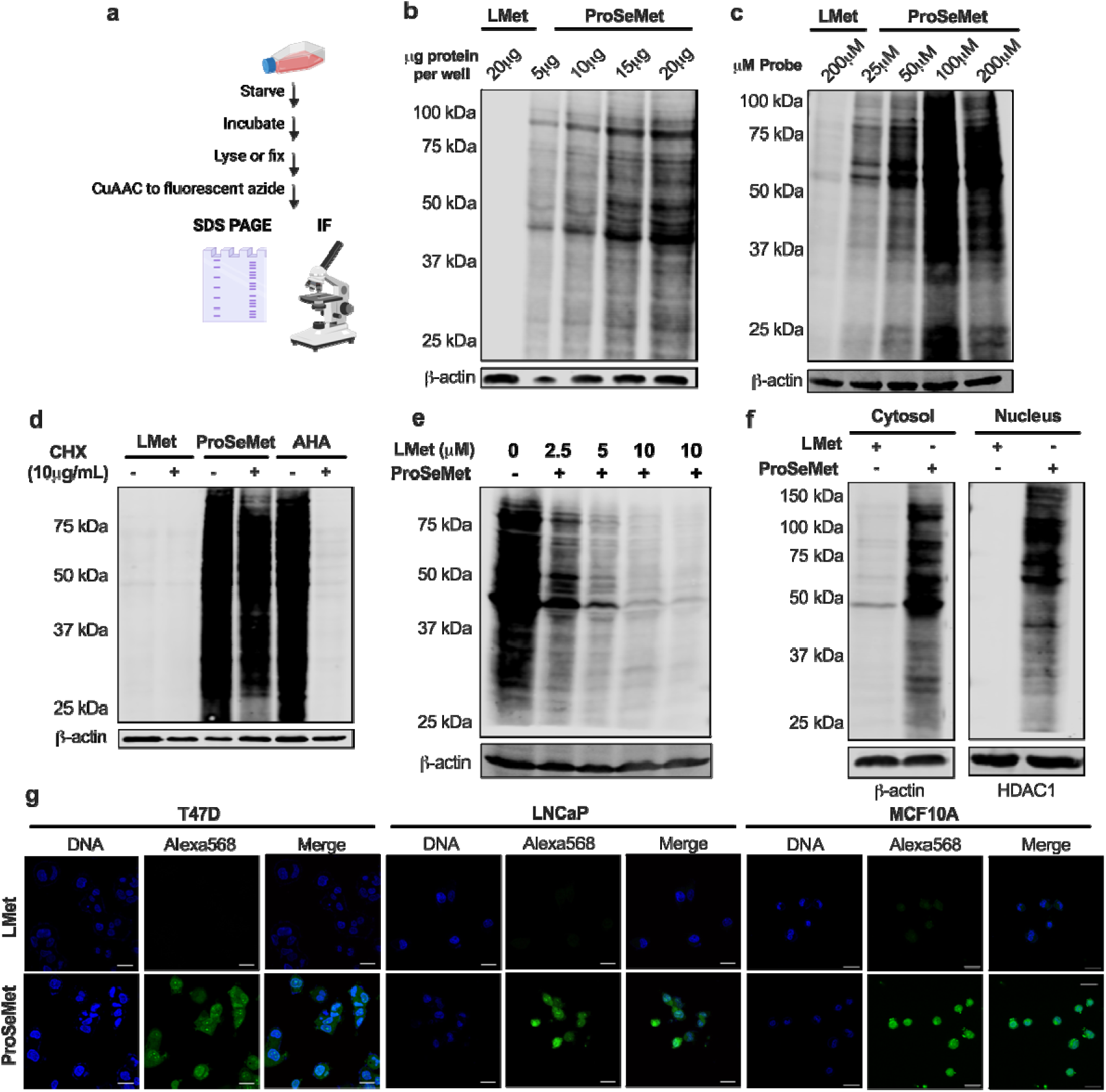
ProSeMet propargylates nuclear and cytosolic proteins in live cells. **a.** Workflow for gel– and IF-based profiling of ProSeMet target proteins. Cells were starved of L-Met by incubating for 30 min in L-Met free media. Cells were then lysed or fixed, subjected to CuAAC to attach a fluorophore, and separated via SDS-PAGE or imaged via confocal microscopy. **b**. T47D cells treated with 100 μM ProSeMet or 100 μM L-Met for 16 h were lysed and subjected to click reaction to attach a fluorescent picolyl azide (680 nm). Increasing protein concentration (5-20 μg) was lo ded into each well and separated by SDS-PAGE (*n*=3). **c**. T47D cells were treated with 200 μM L-Met or increasing concentration (25-200 μM) ProSeMet 16 h. Cellular lysates were subjected to CuAAC and separated via SDS-PAGE. (*n*=3). **d.** T47D cells were treated with 100 μM L-Met, ProSeMet, or AHA in the presence or absence of 10 μg/mL of cycloheximide (CHX). After 16 h, lysates were collected and subjected to CuAAC to attach a fluorescent picolyl azide (L-Met, ProSeMet) or fluorescent alkyne (AHA) then separated by SDS-PAGE (*n*=3). **e**. Competition by increasing concentrations of L-Met during 16 h incubation of T47D cells with 100 μM ProSeMet reduces ProSeMet labeling across molecular weights and in a dose-dependent manner (*n*=3). **f**. Cell fractionation of T47D cells treated with 100 μM ProSeMet or 100 μM L-Met for 16 h. (*n*=3). **g**. T47D, LNCaP, and MCF10A cells were treated with 100 μM ProSeMet or 100 μM L-Met 16 h then fixed, permeabilized, subjected to CuAAC to attach a fluorescent picolyl azide (568 nm, pseudocolored green), and counterstained with Hoechst (blue).

### Propargylated proteins are identified with site specific resolution

To determine whether treatment with ProSeMet would enable the identification of propargylated proteins with site-specific resolution from live cells, the SMARCB1-deficient G401 cell line was treated with ProSeMet or L-Met over a time course of 16 h to 48 h, and cellular lysates were generated, followed by protein digestion and LC-MS/MS. Proteomics analysis of digested lysates identified a total of 376 peptide spectrum matches (PSMs) in a total of 149 proteins, corresponding to 123 (82.5%) unique propargylated proteins in ProSeMet treated samples and 27 PSMs (26 proteins, 17.5%) for L-Met (**Fig. 3a, Supplementary data 1**). We also observed an incubation-time dependent increase in the amount of propargylated peptides (**Fig. 3b**), and amino acids harboring propargylation events including lysine, arginine, and histidine. All instances of propargylated amino acids were observed: lysine mono-, di-, and tri-methylation, arginine mono-, and di-methylation, and histidine monomethylation, with an accumulation of higher and lower-ordered propargylation events for lysine and arginine generally occurring over time (16 h to 48 h) (**Fig. 3c**). The observation of higher-ordered propargylation events with lysine and arginine over time is consistent as the human genome does not encode enzymes known to remove propargyl groups from proteins. Motif enrichment analyses suggest that lysine propargylation occurs around basic residues, arginine propargylation exhibits a slight preference hydrophobic proximal amino acids, and histidine propargylation occurs at aliphatic and/or negatively charged sites (**Fig. 3d).** Identified proteins that harbor propargylated amino acids include proteins previously known to be methylated such as HSPA8, ALDOA, ACTG1, and CALM1 in addition to, and predominantly, novel protein targets that have not previously been defined as methylated, including the lemur tyrosine kinase LMTK3 and the 14-3-3σ adapter protein SFN (**Fig. 3e**). Gene ontology (GO) and pathway-process enrichment analysis of propargylated proteins identified proteins involved in a variety of cellular processes including cellular response to stress, protein folding, cytoskeletal organization, protein methylation, and metabolic processes (**Fig. 3f, Fig. S4, Supplementary data 2**). Taken together, these studies leverage the intracellular chemoenzymatic conversion of ProSeMet into ProSeAM, which is then used as a pseudo-methyl donor to define known and novel amino acid methylation events with site-specific resolution.

**Figure 3.**
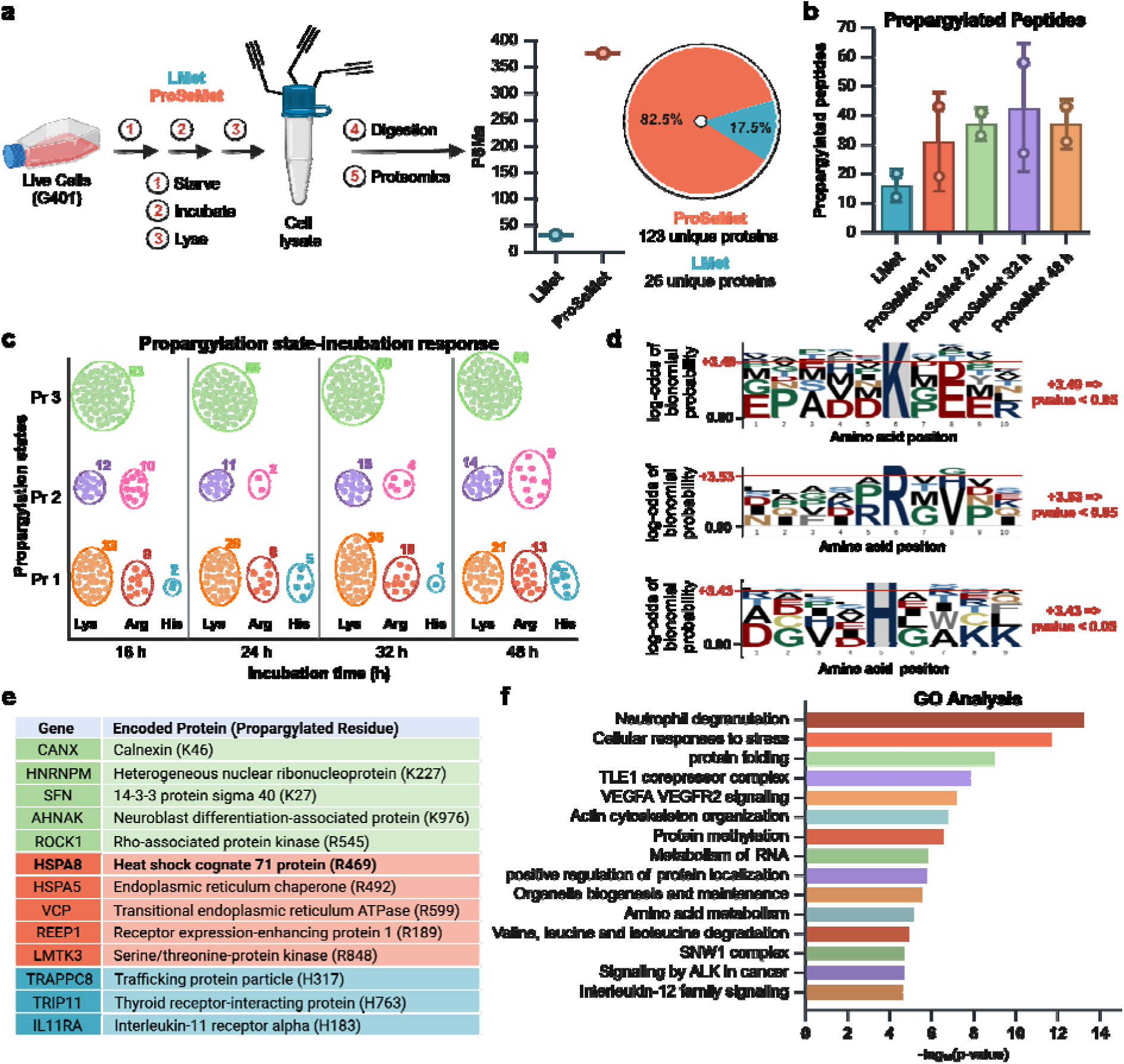
Protein propargylation is mapped with site-specific resolution. **a**. Approach schematic. G401 cells treated with 100 uM ProSeMet with 100 uM L-Met for 16, 24, 36, or 48 h were lysed and processed for LC-MS/MS.. Proteomics analysis identified a total of 376 PSMs corresponding to 149 total proteins. Of these, 123 (82.5%) unique prop rgylated proteins were defined in ProSeMet-treated samples and 27 PSMs (26 proteins, 17.5%) for L-Met. **b.** Total number of propargylated peptides identified via LC-MS/MS across all timepoints, compared to L-Met. **c.** Density plot of peptide propargylation states over the indicated time course. **d.** Sequence motif analysis of propargylated lysine, arginine, and histidine residues. Sequence motif was generated using “probability logo generator for biological sequence motif” plogo v1.2.0 **e.** Representative proteins propargylated in response to ProSeMet incubation and corresponding mapped sites of propargylation. Green, lysine propargylation events; red, arginine propargylation events; blue, histidine propargylation events. **f.** Gene ontology (GO) and pathway-process enrichment analysis of propargylated proteins in response to ProSeMet treatment, (*n=2)*.

### Enrichment of propargylated proteins used to determine breadth of methylproteome

To examine the effect protein enrichment has on mapping the breadth of the methylproteome in live cells, we employed an enrichment-based approach. G401 cells were treated with ProSeMet or L-Met and cellular lysates were subjected to CuAAC to attach a biotin handle, pulled down with streptavidin beads, and analyzed via label-free LC-MS/MS (**Fig. S5**). Using this approach, we identified 707 proteins statistically enriched at *P* values at or below 0.05 and log_2_-fold change greater than 2 (**Fig. S6a**). Of these, 486 proteins have known methylation sites and 221 have uncharacterized methylation sites (**Fig. S7**)^38,39^. Furthermore, 47 proteins were identified at *P* values at or below 0.01 and log_2_-fold change greater than 5 and includes an enrichment of RNA processing proteins (SNRPD3, SNRPD1, SNRPB, SNRPE, QKI), enzymes critical to cell cycle regulation (CDK1), and several methyltransferases (CARM1, KMT2B) (**Fig. S6b**, top 15 genes).

Network analysis of ProSeMet enriched proteins using the GO molecular signatures collection revealed a top list of GO terms involved in biological processes and molecular functions (**Fig. S6c**). Proteins involved in methylation were highly enriched, suggesting pulldown of proteins involved in methyl transfer as well as methyltransferase targets. We further observed positive enrichment of genes associated with p53-mediated signal transduction and DNA damage response (DDR) (**Fig. S6d**). P53 is an important DNA sequence-specific transcription factor known to arrest growth by inducing apoptosis^40^. P53 activity is controlled in part by methylation at multiple key Lys and Arg residues: PRMT5-mediated p53 methylation at R333, R335, and R337 mediate p53 oligomerization and directly affects in DNA-induced apoptosis^41^ whereas competing p53 methylation at K370/K372 by SMYD2/SETD7 can repress or enhance transcription of DDR target genes^7,42,43^. In addition to p53-mediated signaling, we also observed enrichment of proteins involved in RNA binding, processing, and stability, including previously identified PRMT5 targets SNRPD3, SNRPD1, and EEF1A1^44–46^. Our data supports previous studies suggesting heavy methylation of RNA-processing proteins.

Further pathway analysis using the REACTOME gene set identified gene clusters in actin dynamics, cell cycle control, WNT signaling, and transcriptional regulation (**Fig. S8**). Within these gene clusters, we found a series of proteins involved in adhesion dynamics, including the Rho GTPases RhoA, RhoG, and CDC42 and their downstream effectors, such as PAK2. Of these, only RhoA and PAK2 are known to be methylated^38,47^. We further identified key components of the WNT signaling pathway, including G3BP2, a DVL3-associated protein and positive regulator of WNT signaling that is methylated on R432, R438, R452, and R468 as well as WNT5A, which has no previously characterized methylation sites^38,39,48^. Collectively, these experiments identified dozens of known PRMT5, CARM1, EZH2, and other methyltransferase substrates in known pathways as well as novel targets. These pre– and post-enrichment approaches highlight the prevalence of methylation in key signaling pathways and biological processes, and provide a chemical proteomics scaffold to survey the methylproteome in an unbiased manner.

### ProSeMet-mediated chemoproteomics identifies HSPA8 arginine methylation

Label Free Quantification (LFQ) analysis of unenriched propargyl containing proteins identified multiple novel and known sites of arginine, lysine, and histidine methylation (**Fig 3e, Supplementary data 1**) in proteins including the constitutively expressed member of the HSP70 family of heat shock proteins HSPA8, which has essential roles in nascent protein folding as a chaperone protein^49^. Propargylation of HSPA8 was observed at three residues with mono-propargylation of lysine 3 (HSPA8 K3me1), tri-propargylation of lysine 384 (HSPA8 K384me3), and mono-propargylation of arginine 469 (HSPA8 R469me1) (**Fig. 4a, Supplementary data 1, Fig. S9**). HSPA8 arginine propargylation (HSPA8 R469me1) was consistently observed across all timepoints surveyed from 16 h to 48 h (**Fig. 4b**). Among HSP70 proteins, arginine 469 (R469) is conserved and is located within the substrate binding domain, which is only accessible in the ATP-bound, open conformation^50^. Previous studies indicate that this residue undergoes monomethylation catalyzed by CARM1 and PRMT7, thus validating our observed HSPA8 R469 propargylation (**Fig. 4c**)^50,51^. PRMT7-mediated HSP70 R469me1 enhances cytoprotective mechanisms and stress response, consistent with the ability of HSP70 to bind client proteins ^50^. In contrast, CARM1-directed HSP70 R469me1 has been shown to increase gene expression via recruitment of TFIIH during transcription initiation in a manner that is independent of its chaperone activity^51^.

**Figure 4.**
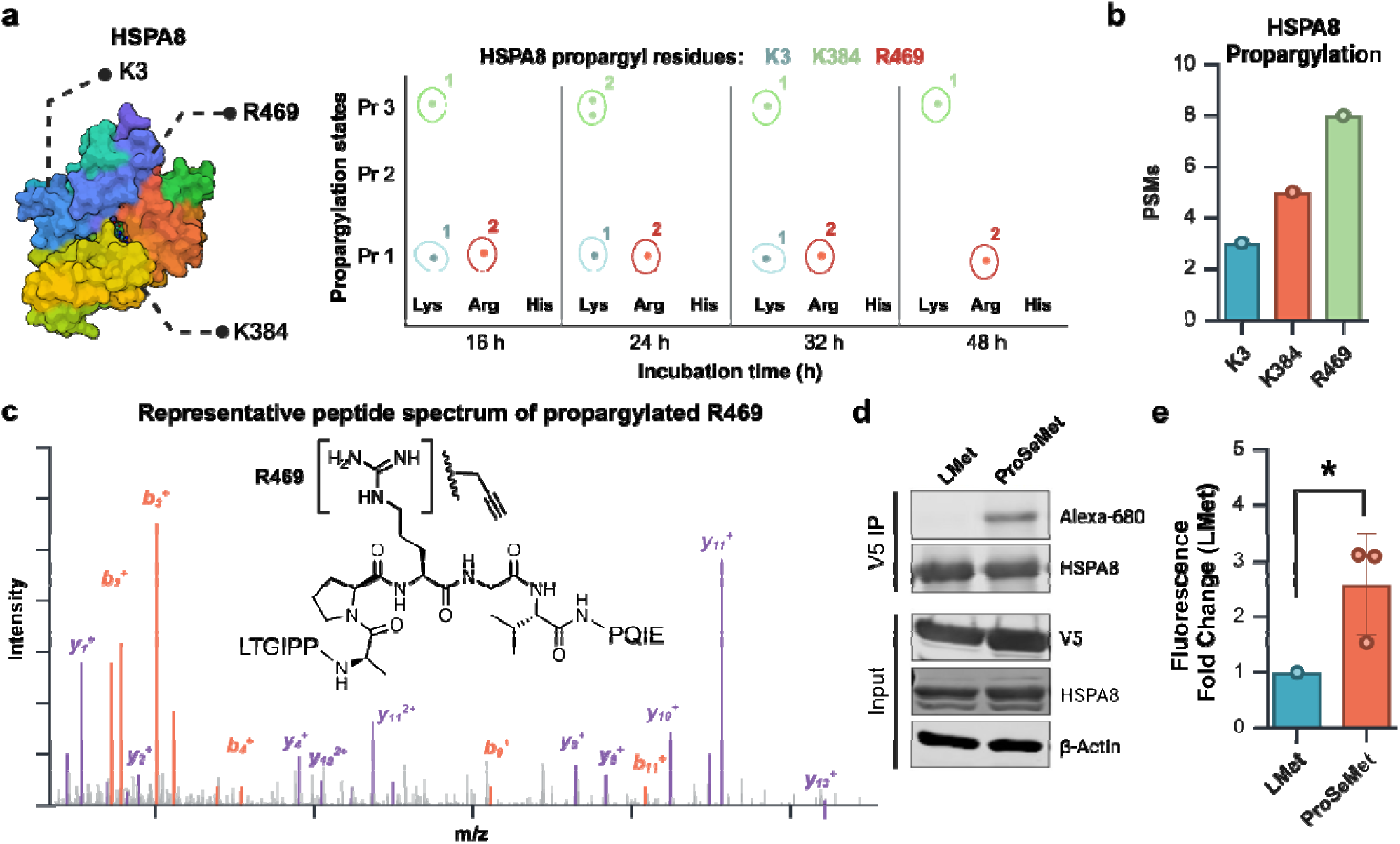
HSPA8 harbors novel propargylation events. **a**. Schematic of HSPA8 propargylation states defined from G401 cells treated with ProSeMet or L-Met for 16, 24, 36, or 48 h, using LC-MS/MS. Propargylation of HSPA8 K3, K383, and R489 were observed across time points. Propargylation sites overlayed on PDB file 2V7Z. **b.** PSM distribution of propargylated HSPA8 sites with 8 PSMs observed for HSPA8 R469me1, over 3 PSMs for HSPA8 K3me1, and HSPA8 K384me3. **c.** Model MS2 peptide spectrum map of HSPA8 mono-propargylated R469. **d.** HSPA8-V5 immunoprecipitation with lysate from HEK293T cells treated with ProSeMet or equimolar L-Met for 24 h, followed by CuAAC to attach a fluorophore azide. **e.** Quantification of (d). (*n=3*), *, p ≤ 0.05.

To empirically determine whether HSPA8 is propargylated, we transiently overexpressed a V5-epitope tagged HSPA8 protein in HEK293T cells and then treated the cells with L-Met or ProSeMet for 24 h, followed by cell lysis. HSPA8 was enriched following V5 immunoprecipitation, after which we used CuAAC to functionalize propargyl groups with a fluorophore-conjugated azide. The resulting proteins were then analyzed via SDS-PAGE and in-gel fluorescence. Fluorescent labeling demonstrates that HSPA8 is selectively propargylated following exposure to ProSeMet (**Fig. 4d, e**). These data demonstrate the utility of our ProSeMet-driven chemoproteomics strategy to uncover existing and novel protein methylation events, thus enabling the identification of methylation associated biological functions and disease related biomarkers.

### ProSeMet-directed propargylation is detectable *in vivo*

The ability of ProSeMet to produce propargylated proteins *in vivo* was explored by administering ProSeMet to a series of model organisms including *A. thaliana, C. elegans, S. cerevisiae, and M. musculus* (**Fig 5a, e)**. ProSeMet or L-Met was administered in growth media to *A. thaliana, C. elegans, or S. cerevisiae* and incubated for up to 48 h, after which lysates were generated and subjected to CuAAC for fluorophore conjugation. For *A. thaliana, C. elegans* and *S. cerevisiae*, treatment with ProSeMet labels proteins across molecular weights (**Fig. 5b-d**). We then administered adult male and female BALB c/J mice ProSeMet. We aimed to assess the impact of existing dietary L-Met on ProSeMet-directed labeling efficiency by dividing mice into the following cohorts: (1) dietary restriction for 12 h prior to L-Met administration; (2) no dietary restriction prior to ProSeMet administration, “fed;” or (3) dietary restriction for 12 h prior to ProSeMet administration, “starved” (**Fig 5e**). In all cohorts, the dietary source of L-Met was restored immediately following L-Met/ProSeMet administration. Mice were administered 15 mg ProSeMet or the equimolar equivalent of L-Met in normal, sterile saline via intraperitoneal (IP) injection. This ProSeMet dose is equivalent to the daily dietary intake of L-Met for an adult mouse^52^. To compare the labeling efficiency between groups, tissue from a diverse array of organs were collected from mice 12 h after ProSeMet or L-Met delivery. The tissues harvested from mice in all conditions were subjected to CuAAC to attach a fluorophore-conjugated azide, then analyzed via fluorescent IHC or lysis followed by SDS-PAGE. Fluorescent labeling across organ systems demonstrates that ProSeMet can diffuse across tissues and penetrate the blood-brain barrier (BBB) (**Fig. 5f-h Fig. S10**). Labeling was detected in both fed mice and mice experiencing dietary restriction to reduce intracellular L-Met abundance prior to ProSeMet administration (**Fig. 5f-h, S10**). These combined data suggest that mice starved prior to ProSeMet injection had increased ProSeMet labeling in most tissues including the brain and lungs, whereas mice fed prior to ProSeMet administration had increased labeling in the heart. Additional SDS-PAGE experiments suggest ProSeMet labeling of the methylproteome occurs as early as 3 h (data not shown). Collectively, these data demonstrate ProSeMet can be utilized in *in vivo* model organisms to propargylate proteins from both diverse model organisms across the phylogenetic tree, as well as diverse organ systems.

**Figure 5.**
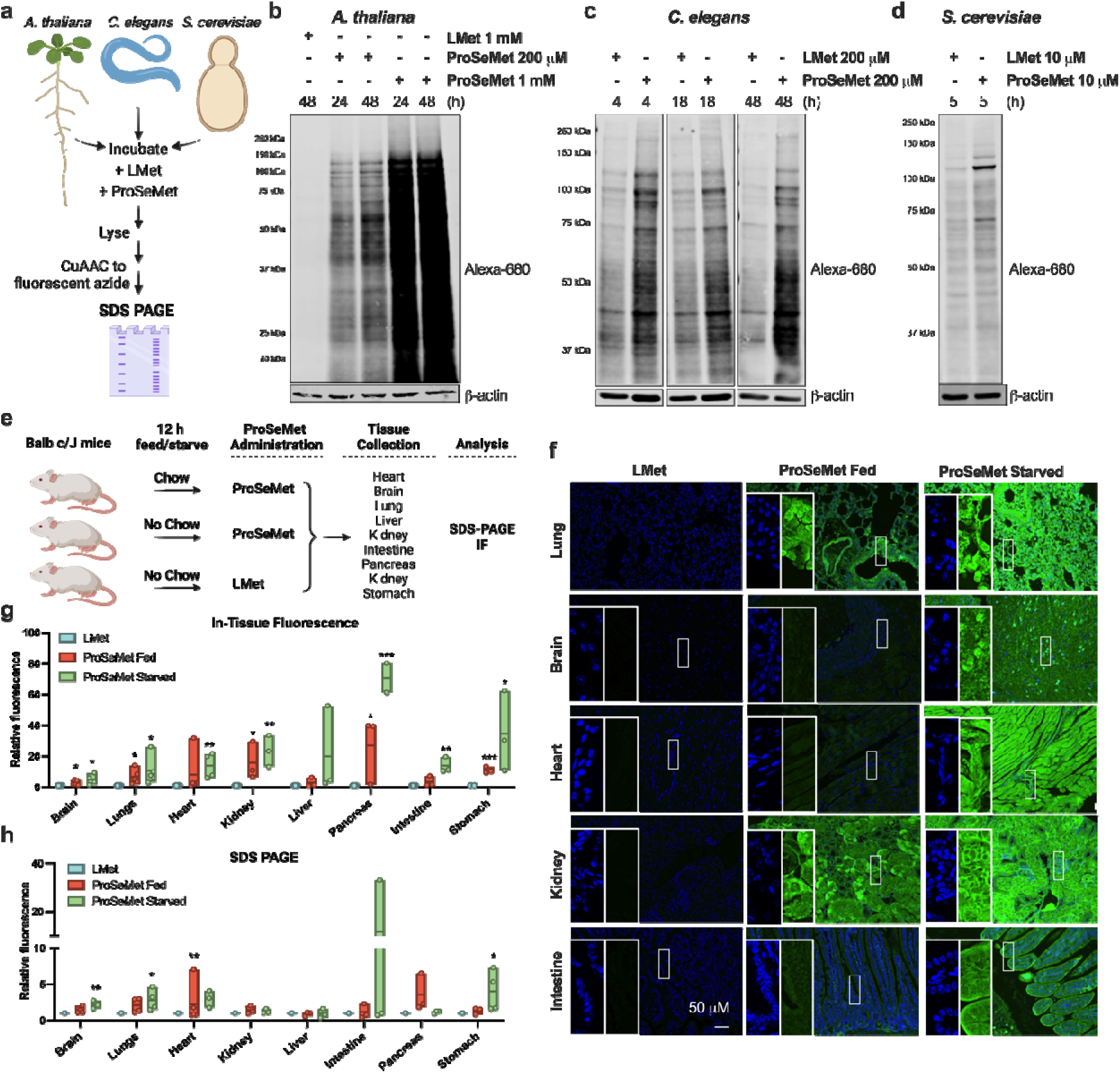
ProSeMet propargylates proteins *in vivo*. **a**. Schematic of propargylation strategy in model organisms. **b.** Lysates extracted from *A. thaliana* root tips from 5-day-old seedlings treated with 200 μM or 1 mM ProSeMet or equimolar L-Met for 24 or 48 h. Lysates were subjected to CuAAC to attach a fluorescent picolyl azide (680 nm), (*n=2*). **c.** Lysates extracted from *C. elegans* treated with 200 μM ProSeMet or equimolar L-Met for 4, 18, or 48 h. Lysates were subjected to CuAAC to attach a fluorescent picolyl azide (680 nm), (*n=2*). **d.** Lysates extracted from *S. cerevisiae* starved for 30 min in Met-Cys-media followed by treatment with 10 μM ProSeMet or equimolar L-Met for 5 h. Lysates were subjected to CuAAC to attach a fluorescent picolyl azide (680 nm), (*n=2*). **e.** Workflow for *in vivo* administration of ProSeMet or L-Met and subsequent analysis of organs for propargylation. **f**. Organs extracted from mice treated with 15 mg ProSeMet or equimolar L-Met via IP injection while being fed or starved for 12 h were fixed in neutral buffered formalin and paraffin embedded. Tissue sections were subjected to CuAAC to attach a fluorescent picolyl azide (568 nm, pseudocolored green) and counterstained with DAPI (blue). In-tissue fluorescence analysis of lung, brain, heart, kidney, and intestine demonstrates successful blood-brain barrier penetrance of ProSeMet, as well as pan-organ labeling. Scalebar represents 50 μM (*n* ≥ 3). **g**. Quantification of in-tissue fluorescence in brain, heart, and lungs (*n* ≥ 3). **h**. Immunoblot densitometry normalized to β-actin and relative to L-Met control (*n* ≥ 3). All tissues show increased protein labeling compared to L-Met; no significant difference observed between mice that were fed or starved prior to ProSeMet administration. *, *p* ≤ 0.05; **, *p* ≤ 0.01*; ***, p* ≤ 0.001.

### *In vivo* protein methylation events are defined with site-specific resolution

To examine whether site-selective arginine, lysine, and histidine propargylation events can be detected *in vivo*, perfused brain, heart, and lung tissues from mice treated with L-Met or ProSeMet as described above (**Fig. 4e**) were digested and processed for LC-MS/MS. Proteomics analyses of digested tissue derived lysates identified an increase in propargyl sites in proteins derived from murine tissue in both the fed and starved groups (see **Fig. 5e** for schematic) when compared to L-Met (**Fig. 6a, Supplementary data 3**). Mice starved prior to ProSeMet administration had increased ProSeMet labeling in the heart (Mean PSM = 29), whereas mice maintained on normal chow prior to ProSeMet administration had increased labeling in the brain (Mean PSM = 25.5) and lungs (Mean PSM = 16). These data are consistent with our bulk analyses of propargylated proteins from in tissue fluorescence and SDS-PAGE (**Fig. 5f-h, Fig S10)**, validating the reproducible nature of leveraging this ProSeMet-driven pseudomethylation strategy for methylome profiling. Network analysis of propargylated proteins using the Gene ontology (GO) and pathway-process enrichment analysis of propargylated proteins identified an enrichment of propargylated proteins involved in organ specific molecular functions, biological processes, and cellular compartmentalization (**Fig. 6b, Fig. S11**, **Supplementary data 4**). Brain-associated propargylated proteins are enriched in protein clusters associated with cellular metabolism and classical neuronal function. In contrast, we observed propargylated proteins associated with structural proteins and muscle related biological processes in the heart, and metabolic and energy-related processes representative of pulmonary function, respectively. Pathway analysis of gene clusters using the KEGG, REACTOME, and WIKIPTHWAYS corroborates the observed tissue-related propargylation, with significant enrichment of neuronal– and metabolic-related pathways in the brain, muscle– and metabolic-related pathways in the heart, and signaling and muscle related pathways in the lung (**Fig. 6b, Fig. S11**). Taken together, these observations highlight the utility of leveraging ProSeMet-driven pseudomethylation for the *in vivo* discovery of methylation events with site-specific resolution.

**Figure 6.**
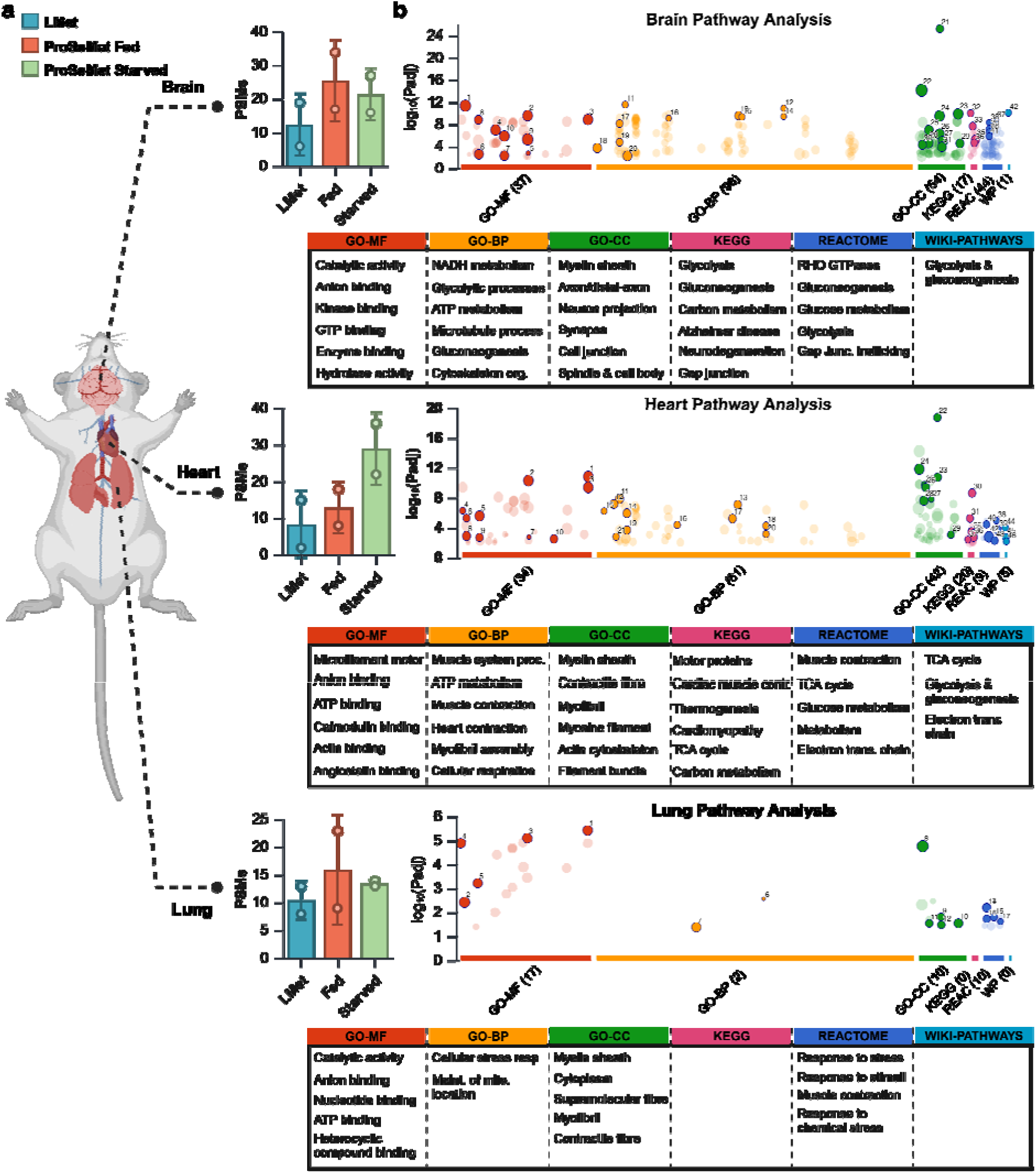
In vivo propargylation is defined with site-specific resolution. **a**. Proteomics analyses of perfused brain, heart and lung tissues from mice treated with L-Met or ProSeMet. Peptide spectral matches (PSMs) of identified propargylated amino acids in ProSeMet fed and starved samples when compared to L-Met control. All samples were filtered for contaminant hemoglobin PSMs. **b.** Gene ontology (GO) and pathway-process enrichment analysis of propargylated proteins, *n=2*.

## Discussion

It is increasingly recognized that non-histone protein methylation is an important component of cell signaling and that comprehensive characterization of the methylproteome will require development of new experimental and analytical pipelines. Herein, we describe the application of the chemical probe ProSeMet, which can be used to propargylate proteins in the natural cellular environment. In cells, ProSeMet is enzymatically converted to ProSeAM by cellular MAT enzymes, which can subsequently be used by diverse methyltransferases in both the cytoplasm and nucleus to deposit a biorthogonal alkyne on target proteins (**Fig. 1**). Our approach extends previous *in situ* methodologies^27–35^ to allow consideration for the biological impetus behind methylation events; this approach allows dynamic, global mapping of the methylproteome.

Using ProSeMet, we developed a reproducible platform for analysis of protein methylation in live cells and in diverse living organisms. For the first time, we demonstrated that ProSeMet-mediated propargylation can be detected using SDS-PAGE and confocal microscopy in cancerous (T47D, LNCaP, G401, 293T) and untransformed (MCF10A) cell lines. Our results indicate that ProSeMet can successfully penetrate the plasma and nuclear membranes to yield labelled proteins across molecular weights (**Fig. 2**). In addition to our *in vitro* results, we report, to our knowledge, the first application of L-Met/SAM analogues in a series of model organisms including *A. thaliana*, *C. elegans*, *S. cerevisiae*, and *M. musculus* (**Fig. 5**), which also suggests diverse applications of ProSeMet, ranging from agriculture to mechanistic studies. To this purpose, ProSeMet administered to Balb c/J mice via IP injection propargylated proteins across tissues including in the heart, lungs, and brain **(Fig. 5**). These data suggest that ProSeMet can be used to study the role of methylation in the pathophysiology of diseases affecting these and potentially other organ systems. We further hypothesize that ProSeMet can be widely leveraged in other *in vitro* and *in vivo* models.

Likewise, we developed two proteomics platforms to identify propargylated proteins, using both native lysate as well as an eluate enriched for propargylated proteins following streptavidin precipitation. Using the SMARCB1-deficient cell line G401, we treated cells with ProSeMet or L-Met, generated cell lysate, and processed this lysate for LC-MS/MS in the absence of functionalization and enrichment. This approach identified a series of both known and novel protein methylation events, including the heat shock chaperone protein HSPA8 (**Fig. 4**). Some proteins within the heat shock family are ubiquitously expressed at relatively high intracellular concentrations^53^, which may explain the reproducible detection of HSPA8 R469me1 in our LC-MS/MS runs. PRMT9-mediated HSPA8 Arg (R76 and R100) methylation has been implicated in hepatocellular carcinoma^54^, and CARM1 and PRMT7 have been shown to methylate HSP70 proteins including HSPA8 on R469 in a chaperone/client independent– and dependent-manner, respectively^50,51^. Our studies presented here demonstrate robust HSPA8 R469 methylation, and provide an experimental platform for future studies examining the cellular contexts of PRMT-mediated HSPA8 methylation, which when coupled with mechanistic and functional studies, are likely to provide further insight into the biological function(s) of HSPA8 R469me1. In addition to HSPA8 Arg methylation, this unenriched approach defined all species of canonical protein methylation, including Arg, His, and Lys, as well as mono-, di-, and tri-methylation. The detection of Lys tri-propargylation was unexpected; we hypothesized this propargylated species may be rare as steric hindrance of the prior two deposited propargyl groups could render the tri-propargylation event sterically challenging for methyltransferases responsible for tri-propargyl deposition (**Fig. 1c)**.

We also observed labeling in a variety of model organisms, demonstrating that this ProSeMet-driven chemoenzymatic approach may be amenable to use in a variety of living organisms, including and beyond those tested here. Leveraging model organisms for these studies is especially attractive given the availability of a wide variety of genetic mutant strains to examine the mechanisms of methylation, as well as the benefit that model organisms such as *S. cerevisiae* encode a smaller array of methyltransferases within their genome. The latter may render these organisms as powerful tools to unmask and map both the methylproteome as well as the network of responsible methyltransferases, which is already known to have implications in disease etiologies such as cancer. The utility of complex model organisms such as murine models cannot be underscored – the observation that ProSeMet labels proteins across the blood-brain barrier as well as in all organ systems examined suggests that ProSeMet labeling in *in vivo* murine models may have applications in studying tissue-specific methylation in complex disease states such as cancer or neurological disorders. Collectively, these suggest broad applicability in addressing both unbiased and biased questions related to cellular methylation events in *in vitro* and *in vivo*.

Using G401 cells, we then adapted our pipeline and leveraged biotin conjugation and streptavidin enrichment to define non-histone proteins known to be methylated by PRMT1 (NONO)^55^, PRMT5 (SNRPD3, SNRPD1)^45^, CARM1 (SNRPB, SF3B4)^7^, and EZH2 (STAT3, PCNA)^11,56^ (**Fig. S6**. We also identified the catalytic subunits (PPP2CA, PPP4C, PPP6C) of several protein phosphatases known to be methylated by the leucine carboxyl methyltransferase LCMT1^57^. In addition to these and other known methyltransferase targets, we identified 221 proteins with uncharacterized methylation sites with unknown biological function (**Fig. S7**). Our enrichment proteomics data also indicated enrichment of the SAM synthetase enzymes MAT2A/B, which are responsible for the coupling of ATP and L-Met to produce SAM^58^. While we cannot discount the possibility that these enzymes were identified because of ProSeAM present in their binding pockets, MAT2A is purportedly methylated on a critical Arg residue in the binding pocket (R264) and this methylation event may serve as a biological switch to control cellular SAM levels^59^. Similarly, our MS data indicates enrichment of several methyltransferases, including the Lys methyltransferases KMT2A/B, the Arg methyltransferases PRMT1, 3, 5 and CARM1, and the RNA methyltransferase NSUN2, all of which are known to be methylated^38^. To expand upon the approaches taken here, future chemoproteomics analyses of ProSeMet targets could focus on elucidation of the molecular mechanisms by which methylation of methyltransferases controls their function.

The ProSeMet pipelines described herein allows profiling of methyltransferase targets *in vitro* and *in vivo* while allowing a comprehensive survey of canonical and non-canonical methylated residues. Our approach has several advantages over previous methodologies: ProSeMet readily enters live cells and whole organisms without toxicity; ProSeMet-mediated propargylation considers the biological impetus for a methylation event; ProSeMet and the corresponding SAM analogue ProSeAM can be used by native methyltransferases in cell lines and *in vivo*; and the CuAAC reaction allows for identification of propargylated proteins, with or without enrichment, via LC-MS/MS. This platform provides a new avenue to dissect the methylation-controlled crosstalk and signal transduction responsible for cellular function.

## Methods

### Synthesis of ProSeMet

Propargyl-Selenium L-Methionine (ProSeMet) was synthesized follow a previously described protocol^60^ with modified purification method. L-selenohomocystine (200 mg, 0.55 mmol) was dissolved in dry ethanol (25 mL), NaBH_4_ (126 mg, 3.30 mmol) was added under nitrogen, and the reaction mixture was stirred at room temperature (RT) for 15 min. NaHCO_3_ (152 mg, 3.3 mmol) was added to the reaction mixture in one portion, followed by propargyl bromide (250 µL, 3.3 mmol), and the resultant reaction mixture was stirred RT for 12 h. The solvent was removed under reduced pressure, and the crude compound was dissolved in distilled water (6 mL) and 1% TFA (60 µL). The resulting solution was directly loaded to neutral alumina, and then 50% EtOAc and hexane (50 mL) were used to wash the nonpolar impurities, followed by 50% methanol and DCM (50 mL) to remove polar impurities. ProSeMet was eluted by NH_4_OH solution (50 mL) from neutral alumina column, which was concentrated and lyophilized to afford the pure product (92 mg, 76%).

**ESI-MS**: Expected mass for C_7_H_11_NO_2_SeK^+^: 259.24 [M + K]^+^; found: 259.815. (**Fig. S12**).

**^1^H NMR**: ^1^H NMR (600 MHz, D_2_O) δ 3.84 – 3.81 (m, 1H), 3.30 (s, 2H), 2.84 (t, *J* = 8.0 Hz, 2H), 2.64 (s, 1H), 2.30 –2.20 (m, 2H) (**Fig. S13**).

**^13^C NMR**: (151 MHz, D_2_O) δ 175.29, 54.90, 54.64, 31.67, 19.30, 19.09, 6.31. (**Fig. S14**).

### Cell Culture

Cells were purchased from ATCC and maintained at 37 °C and 5% CO_2_. HEK293T, T47D, LNCaP, and G401 cells were cultured in either DMEM or RPMI supplemented with 10% (V/V) fetal bovine serum (FBS) and 1% (V/V) penicillin/streptomycin (pen/strep) (100 µg/mL). MCF10A cells were cultured in DMEM/F-12 supplemented with 3% (V/V) FBS, cholera toxin (100 ng/mL), EGF (20 ng/mL), hydrocortisone (0.5 mg/mL), insulin (10 µg/mL), and 1% (V/V) pen/strep. For LMet depletion experiments, cells were cultured in cysteine– and methionine-free DMEM (Gibco 21013024) supplemented with pen/strep, L-Cystine 2HCl (63 mg/mL), and dialyzed serum (Gibco A3382001). Cycloheximide (CHX) was purchased from Sigma (01810).

#### Cell and Organism Lysis and Immunoblotting

*H. sapiens* whole cell lysate was generated by lysing cells on ice in RIPA buffer (50 mM TrisHCl [pH 8], 150 mM NaCl, 1% NP-40, 0.5% sodium deoxycholate, 0.1% SDS) supplemented with protease and phosphatase inhibitors. *A. thaliana* whole cell lysate was generated by grinding 1 g of root tissue to a powder in liquid N_2_ in a mortar and pestle and then lysed with 1 mL of RIPA buffer with further grinding. The suspension was then filtered through a 70 μM cell strainer and centrifuged at 1200 x *g* for 5 min at 4°C., and the supernatant obtained after centrifugation was used as protein lysate. *C. elegans* whole organism lysate was generated by lysing nematodes in RIPA buffer, including 25 cycles of douncing. Lysates were centrifuged at 16,000 *x* g for 10 min at 4°C, and soluble lysate collected. *S. cerevisiae* whole cell lysate was generated by lysing cells in RIPA buffer. Lysates were centrifuged 6,500 *x* g, 10 min at 4°C, and soluble lysate collected. Proteins were separated using 12 or 16% SDS-PAGE. When applicable, proteins were fluorophore labeled via CuAAC. Proteins were then transferred to nitrocellulose membranes and visualized directly via fluorophore or visualized after primary antibody incubation (Odyssey, Li-Cor). Primary antibodies used were as follows: β-actin (Millipore-Sigma MAB1501), V5 (CST80076S), HSPA8 (CST8444S), and HDAC1 (Upstate 05-614).

#### Copper-catalyzed Azide-Alkyne Cycloaddition (CuAAC) reaction

CuAAC on whole cell lysate, fixed cells, or tissue sections was performed using 1 mM CuSO4, 1.5 mM THPTA, 3 mM sodium ascorbate (NaAsc) and indicated concentration of azide (**Table S1**). All azides and alkynes were purchased from ClickChemistryTools. For whole cell lysate, CuAAC reagents were added directly to lysate, incubated for 30 or 60 m at RT, then prepared for downstream analysis. For confocal microscopy, cells were fixed in 3.7% PFA and permeabilized with 0.5% Triton-X. CuAAC reagents were added directly to cells, then incubated, rocking, for 45 m. For IHC, tissue sections were permeabilized as described below and CuAAC reagents supplemented with 5% DMSO and 0.2% TX-100 were added directly to tissue section for 1 h prior to proceeding with the confocal microscopy pipeline described below.

#### Biotin-Streptavidin Pulldown

G401 cells were incubated in L-Met free media for 30 min, treated with 100 µM ProSeMet or L-Met for 16 h, then lysed with RIPA buffer. Protein lysate (1 mg/IP) was diluted to 1 mL final volume in PBS, then subjected to CuAAC for 1 h at RT to attach an azide conjugated biotin moiety (ClickChemistryTools). After CuAAC, protein was precipitated in acetone. Briefly, 4 volumes of ice-cold acetone were added to the post-CuAAC lysate, followed by incubation at –20 °C for 1 h. Precipitated protein were centrifuged at 9,000 *x* g for 10 min and resuspended in PBS with 5% SDS. After resuspension, SDS solution was diluted to 0.5% in PBS, then magnetic streptavidin beads (Pierce) were added directly to the resuspended protein. Protein and beads were incubated for 2 h at RT, rocking, and then beads were washed 3X with ice-cold RIPA buffer and 3X with ice-cold PBS. Beads with attached proteins were stored at –80 °C until LC-MS/MS analysis.

#### Immunoprecipitation

HEK293T cells were transiently transfected with plasmid HSPA8_pLX307 (Addgene Plasmid #98343). 48 h post transfection, cells were washed twice in 1X PBS, and incubated in Cys-Met-media (Invitrogen) for 30 min and then treated with 200µM ProSeMet or L-Met for 24 h. At 72 h post-transfection, cells were lysed with RIPA buffer. Protein lysate (4.5mg) was diluted in IP Buffer (20 mM Tris pH 7.5, 150 mM NaCl, 5 mM MgCl_2_, 1% NP-40) for immunoprecipitation (IP) with 0.84 µg V5 antibody (CST80076) and 40 µL washed Protein G beads (Invitrogen). Post IP, beads were washed in IP Buffer, followed by PBS, and resuspended in 1X LDS before boiling for 5 min at 95°C. Denatured proteins were loaded into centrifugal 30 kDa filter (Millipore), and diluted with 400 µL of DI H2O. Filters were spun at 10,000 RPM for 10 min, 200 µL of H2O was added, and spun at 10,000 RPM for 10 min. Remaining volume containing proteins was subjected to CuAAC for 30 min at RT to attach an azide conjugated fluorophore. After CuAAC, proteins were loaded into a centrifugal 30 kDa filter, diluted with 300µL of H2O, and spun for 10 min at 10,000 RPM. 200 µL of additional H2O was added and filter was spun at 10,000 RPM for 25 min. The remaining solution containing proteins was separated using SDS-PAGE, and immunoblots were performed as described above.

#### Annexin V/PI Staining

T47D cells were incubated with 5 µM or 20 µM of ProSeMet for 24 h, detached with trypsin (Biolegend), and stained using Annexin V/PI following manufacturer’s protocol (Biolegend). Cells were analyzed via flow cytometer (BDFACSymphony A3) within 1 h.

#### Confocal microscopy

Live cells were plated and after 16 h, cells were washed to remove excess L-Met and incubated in methionine-free supplemented DMEM (Gibco) for 30 min to remove endogenous methionine. Subsequently, 100 µM or 200 µM concentrations of ProSeMet was added to the cells. After 16 h, cells were fixed with 4% PFA and permeabilized using 0.5% Triton X-100. The CuAAC reaction was then performed directly on the fixed and permeabilized cells as described to attach a 488 or 568 nm picolyl azide-conjugated fluorophore (ClickChemistryTools) and nuclei were counterstained using Hoechst. Cells were imaged on Leica SP8 confocal microscope and images were processed and analyzed using ImageJ and Python.

## Mass spectrometry

### Mass spectrometry for unenriched samples

#### Sample Preparation

ProSeMet and L-Met modified lysates were digested using EasyPep™ Mini MS Sample Prep Kit (Thermo Scientific). 100 µg of proteins were treated with 50 µL of reduction and 50 µL of alkylation buffer and then incubated at 95°C for 10 min in the dark. Reduced and alkylated protein samples were cooled to RT and digested with a digestion cocktail of 50 µL (trypsin and Lys-C) or 50 µL of (Glu-C) for 24 h. Resulting peptides were desalted with EasyPep™ Mini MS peptide cleanup column (Thermo Scientific) and dried under vacuum.

#### LC-MS/MS for Unenriched samples

Digested samples were resuspended in Buffer A (0.1% FA in water) and the peptide amount was determined by Pierce™ Quantitative Peptide Assays & Standards (Thermo Fisher Scientific) according to manufactures instructions. Samples were injected into a nanoElute UPLC autosampler (Bruker Daltonics) coupled to a timsTOF Pro2 mass-spectrometer (Bruker Daltonics). The peptides were loaded on a 25 cm Aurora ultimate CSI C18 column (IonOpticks) and chromatographic separation was achieved using a linear gradient starting with a flow rate of 250 nl/min from 2% Buffer B (0.1% FA in MeCN) and increasing to 13% in 42 min, followed by an increase to 23% B in 65 min, 30% B in 70 min, then the flow rate was increased to 300 nl/min and 80% B in 85 min, this was kept for 5 min. The mass-spectrometer operated in positive polarity for data collection using a data-dependent acquisition (ddaPASEF) mode. The cycle time was 1.17 s and consisted of one full scan followed by 10 PASEF/MSMS scans. Precursors with intensity of over 2500 (arbitrary units) were picked for fragmentation and precursors over the target value of 20,000 were dynamically excluded for 1 min. Precursors below 700 Da were isolated with a 2 Th window and ones above with 3 Th. All spectra were acquired within an m/z range of 100 to 1700 and fragmentation energy was set to 20 eV at 0.6 1/K0 and 59 eV at 1.60 1/K0.

#### Database search (MSFragger)

MS raw files were searched FragPipe GUI version 20 with MSFragger (version 3.8) as the search algorithm. Protein identification was performed with the human Swisspot database (20′456 entries) with acetylation (N-terminus), and oxidation on methionine was set variable modification. To account for the mass shift introduced by the propargyl handle, a variable mass shift of 38.0157 Da on Lysine, Arginine, Histidine with a maximal occurrence of 3 propargyl respectively, was set. Carbamidomethylation of cysteine residues was considered a fixed modification. Trypsin or GluC was set as the enzyme with up to two missed cleavages. The peptide length was set to 7–50, and the peptide mass range of 500–5000 Da. For MS2-based experiments, the precursor tolerance was set to 20 ppm and fragment tolerance to 20 ppm. Peptide spectrum matches were adjusted to a 1% false discovery rate using Percolator as part of the Philosopher toolkit (v5). For label-free quantification, match-between-runs were enabled. All downstream analysis was performed in R (version 2023.03.0). Individual samples were normalized to the mean of all quantified peptides.

#### Data analysis

For the unenriched pipeline, following database search with MSFragger, contaminant proteins were filtered and PSMs with zero intensities and Match type of “unmatched” were removed. Blood-related hemoglobin PSMs were filtered from mice peptide data. For Gene Ontology (GO) analysis of G401 cells, gene list of propargylated proteins were utilized as input in metascape, with input and analysis species set to *H. sapiens*. For GO analysis of the murine model, a gene list of propargylated proteins was utilized as input in metascape^61^ and gProfiler, with input and analysis species set to *M. Musculus.* Pathway and process enrichment analysis was carried out with the following ontology sources: KEGG Pathway, GO Biological Processes, Reactome Gene Sets, Canonical Pathways, CORUM, WikiPathways, and PANTHER Pathway. All genes in the human and mice genome were used as the enrichment background. Terms with a p-value < 0.01, a minimum count of 3, and an enrichment factor > 1.5 were utilized. p-values were calculated based on the cumulative hypergeometric distribution, and q-values are calculated using the Benjamini-Hochberg procedure. To identify the sequence motif of ProSeMet modified sites, “probability logo generator for biological sequence motif “ plogo v1.2.0 was utilized. Sequences containing 5 residues from the left and 4 residues from the right of modified lysine and arginine sites were utilized, with lysine or arginine as the fixed positions with a p-value <0.05. For sequence motif analysis of histidine, sequences containing 4 residues from the left and 4 residues from the right of modified histidine sites were utilized.

### Mass spectrometry for enriched samples

#### Sample Preparation

On-bead digestion was performed as previously described^62^. To the beads, digestion buffer (50 mM NH_4_HCO_3_) was added, and the mixture was then treated with 1 mM dithiothreitol (DTT) at RT for 30 min, followed by 5 mM iodoacetimide (IAA) at RT for 30 min in the dark. Proteins were digested with 2 µg of lysyl endopeptidase (Wako) at RT overnight and were further digested overnight with 2 µg trypsin (Promega) at RT. The resulting peptides were desalted with HLB column (Waters) and were dried under vacuum.

#### LC-MS/MS for enriched samples

The data acquisition by LC-MS/MS was adapted from a published procedure^63^. Derived peptides were resuspended in the loading buffer (0.1% trifluoroacetic acid, TFA) and were separated on a Water’s Charged Surface Hybrid (CSH) column (150 µm internal diameter (ID) x 15 cm; particle size: 1.7 µm). The samples were run on an EVOSEP liquid chromatography system using the 15 samples per day preset gradient (88 min) and were monitored on a Q-Exactive Plus Hybrid Quadrupole-Orbitrap Mass Spectrometer (ThermoFisher Scientific). The mass spectrometer cycle was programmed to collect one full MS scan followed by 20 data dependent MS/MS scans. The MS scans (400-1600 m/z range, 3 x 10^6^ AGC target, 100 ms maximum ion time) were collected at a resolution of 70,000 at m/z 200 in profile mode. The HCD MS/MS spectra (1.6 m/z isolation width, 28% collision energy, 1 x 10^5^ AGC target, 100 ms maximum ion time) were acquired at a resolution of 17,500 at m/z 200. Dynamic exclusion was set to exclude previously sequenced precursor ions for 30 seconds. Precursor ions with +1, and +7, +8 or higher charge states were excluded from sequencing.

#### MaxQuant

Relative intensity values were calculated from raw data using MaxQuant. Label-free quantification analysis was adapted from a published procedure^63^. Spectra were searched using the search engine Andromeda, integrated into MaxQuant, against Human Uniprot/Swiss-Prot database (20,379 target sequences). Methionine oxidation (+15.9949 Da), asparagine and glutamine deamidation (+0.9840 Da), and protein N-terminal acetylation (+42.0106 Da) were variable modifications (up to 5 allowed per peptide); cysteine was assigned as a fixed carbamidomethyl modification (+57.0215 Da). Only fully tryptic peptides were considered with up to 2 missed cleavages in the database search. A precursor mass tolerance of ±20 ppm was applied prior to mass accuracy calibration and ±4.5 ppm after internal MaxQuant calibration. Other search settings included a maximum peptide mass of 6,000 Da, a minimum peptide length of 6 residues, 0.05 Da tolerance for orbitrap and 0.6 Da tolerance for ion trap MS/MS scans. The false discovery rate (FDR) for peptide spectral matches, proteins, and site decoy fraction were all set to 1 percent. Quantification settings were as follows: re-quantify with a second peak finding attempt after protein identification has completed; match MS1 peaks between runs; a 0.7 min retention time match window was used after an alignment function was found with a 20-minute RT search space. Quantitation of proteins was performed using summed peptide intensities given by MaxQuant. The quantitation method only considered razor plus unique peptides for protein level quantitation.

#### Data analysis

For the enrichment pipeline, following MaxQuant, all data was log2-transformed before further analysis. Contaminant proteins were filtered and missing values were imputed using a normal distribution in Perseus. Differential protein expression was performed on normalized intensity values using the DEqMS R package (https://www.bioconductor.org/packages/release/bioc/html/DEqMS.html)^64^. Samples with a high level of variance calculated using PCA were removed (**Fig. S7**). GSEA was performed on enriched proteins (LFC > 1) using the fgsea R package (https://bioconductor.org/packages/release/bioc/html/fgsea.html). Datasets with log_2_-transformed fold change were analyzed using C5 (GO) and C2 (REACTOME) gene set collections in the Molecular Signatures Database (MSigDB v 7.5.1). Identification of proteins with known or unknown methylation sites was performed by cross referencing the PhosphoSitePlus and ActiveDriver DB databases^65,66^.

### In vivo studies

#### C. elegans

N2: wild-type (Bristol isolate) *Caenorhabditis elegans* were cultured at 20°C on 60 mm nematode growth media (NGM) agar plates with OP50 bacteria grown in Luria Broth (LB). Mixed stage worms were washed into liquid OP50 grown in LB and maintained by shaking at moderate speed at 20°C for 10 d, refreshing the OP50 bacteria at 5 d. The resulting population was washed by allowing the worms to sink via gravity and replacing the M9 buffer (22 mM KH_2_PO_4_, 42 mM Na_2_HPO_4_, 86 mM NaCl, and 1 mM MgSO_4_) approximately 5 times or until liquid ran clear. Worms were then starved by shaking in M9 buffer without bacteria at 20°C for 4 h, pelleted, then added to fresh OP50 with either L-Met or ProSeMet at a final concentration of 200uM. Worms were collected at 4 h, 18 h, and 48 h after starvation was ended by pelleting and washing with M9 buffer approximately 5 times or until liquid ran clear. A final volume of 500uL M9 containing whole worms was flash frozen with liquid nitrogen and stored at –80°C until lysis.

#### A. thaliana

Col-0 ecotype seeds were sterilized and sown on 1/2 Murashige and Skoog (MS) medium supplemented with 1% (w/v) sucrose. The seedlings were grown on plates in the vertical orientation in controlled growth chamber under 16 h light/8 h dark cycles, at a temperature of 20°C, and light intensity of 200 μmol/m^2^/s for 5 d after germination. After 5 days, seedlings were treated with 10 mL of MS media containing either 200 µM or 1 mM ProSeMet or L-Met by pouring the liquid onto the plates and swirling. Plates were then returned to the growth chamber and kept in a horizontal orientation for 24 h or 48 h. Whole roots from the treated seedlings were then harvested and immediately frozen in liquid N_2_ for protein extraction.

#### S. cerevisiae culturing

S288C cells (MATa; ura3-52; leu2Δ1; HIS3; trp1Δ63) were grown in complete YEPD medium with overnight shaking at 30°C. Cells were then diluted to OD = 0.1 in Met-Cys-medium, starved for 30 m, and L-Met or ProSeMet was then added at 10 µM concentrations. When the cells reached OD=1 (about 6.5 h), pellets were flash frozen. Pellets were resuspended in RIPA buffer (50 mM Tris HCl pH8.0, 150 mM NaCl, 1% NP-40, 0.5% Na deoxycholate, 0.1% SDS) supplemented with protease inhibitors. Cells were lysed by beating with 0.3 mg acid washed glass beads for 1 min with 2 min rest on ice, 4 times. Lysates were clarified via centrifugation at 14,000 rpm at 4°C for 10 m.

#### M. musculus

Male and female BALB c/J mice (Jackson labs) were maintained on a standard chow diet. 12 h prior to L-Met or ProSeMet delivery, mice were either starved or maintained on the standard chow diet, after which they were administered 15 mg ProSeMet or equimolar L-Met, resuspended in sterile 0.9N saline, via intraperitoneal (IP) injection. At the time of ProSeMet or L-Met administration, standard chow was returned. Mice were monitored hourly post administration and euthanized after 12 h. All tissue collected was isolated and prepared within 1 h following euthanasia. All mouse experiments were conducted in accordance with protocols approved by the Institutional Animal Care and Use Committee (IACUC) of Emory University School of Medicine (EUCM).

### In tissue fluorescence

Organs extracted from mice treated with ProSeMet or L-Met were immediately placed in 4% formaldehyde solution and left at RT for 24 h. Subsequently, organs were paraffin embedded and sectioned. Tissue slides were rehydrated by washing 3 times with xylenes (mixed isomers), twice with 100% EtOH, once with 95% EtOH, once with 80% EtOH, once with 70% EtOH, and once with PBS. All washes were for 5 m. After the final wash, tissue sections were permeabilized using 0.5% TX-100 in PBS for 1 h. After permeabilization, CuAAC was performed directly on slides using 1 mM CuSO4, 1.5 mM THPTA, 3 mM NaAsc, 25 µM picolyl azide conjugated to a 568 nm fluorophore (ClickChemistryTools), 5% DMSO, and 0.2% TX-100, for 1 h at RT. After reaction completed, slides were washed for 30 min in PBS supplemented with 5% DMSO and 0.2% TX-100, then for 30 min in PBS. Slides were then stained with DAPI (300 nM) for 5 min at RT. Prior to mounting, slides were dehydrated by washing once with 70% EtOH, once with 80% EtOH, twice with 100% EtOH, and twice with xylenes (mixed isomers). All washes were for 5 m. After dehydration, slides were mounted and imaged using a confocal microscope (Leica SP8).

## Supporting Information

Additional experimental details and images, and ^1^H NMR spectra for all compounds (PDF).

## Acknowledgements and funding sources

The authors thank Dr. David Lynn, Dr. Andrew Hong, and the Spangle, Raj, and Hong labs for helpful project discussions. This study was supported by funds from Winship Cancer Institute of Emory University and the NIH (1R35GM150587), to JMS. M.R. acknowledges funding from NIH (1R35GM133719-01). The *S. cerevisiae* work performed by C.Y.J. was funded by the NIH (K12 GM000680, to C.Y.J.). The *C. elegans* work performed by M.R. (T32GM008490-21) was funded by a grant to D.J.K. (NSF IOS1354998). This study was supported in part by the Emory University Cancer Tissue Pathology Core and Integrated Proteomics Core facility (RRID:SCR_023530), both of which are shared resources of Winship Cancer Institute of Emory University and the NIH/NCI (P30CA138292). The indicated figures were created with BioRender. [Fig. 3 (License PF26RS7R7U), Fig. 4 (License ZG26T7IHR3), Fig. 5a (License WJ26RQFQ1T), Fig. 5e (License NP26RQN94I), and Fig. 6 (License PK26RS9KSL).

## Author Contributions

J.F., B.E., M.R., and J.M.S. designed the research. J.F., B.E., R.S.L., R.B.J., C.M.B., A.K.V., C.Y.J., M.F., M.R., K.S., K.K.P., and P.B. performed experiments and analyzed the results. J.M.S., M.R., R.B.D., A.H.C, D.J.K., and D.E.G. supervised the studies. J.F., B.E., and J.M.S. wrote the manuscript with input from all authors.

## Data availability statement

The data generated in this study are publicly available in synapse (in progress).

Correspondence and requests for materials should be addressed to: Dr. Jennifer M. Spangle, Ph.D.

## References

1 Murn, J. & Shi, Y. The winding path of protein methylation research: milestones and new frontiers. Nat Rev Mol Cell Biol 18, 517–527 (2017). 10.1038/nrm.2017.35

2 Michalak, E. M., Burr, M. L., Bannister, A. J. & Dawson, M. A. The roles of DNA, RNA and histone methylation in ageing and cancer. Nat Rev Mol Cell Biol 20, 573–589 (2019). 10.1038/s41580-019-0143-1

3 Fontecave, M., Atta, M. & Mulliez, E. S-adenosylmethionine: nothing goes to waste. Trends Biochem Sci 29, 243–249 (2004). 10.1016/j.tibs.2004.03.007

4 Bedford, M. T. Arginine methylation at a glance. Journal of cell science 120, 4243–4246 (2007).

5 Bannister, A. J. & Kouzarides, T. Regulation of chromatin by histone modifications. Cell Res 21, 381–395 (2011). 10.1038/cr.2011.22

6 Wu, Z., Connolly, J. & Biggar, K. K. Beyond histones – the expanding roles of protein lysine methylation. FEBS J 284, 2732–2744 (2017). 10.1111/febs.14056

7 Biggar, K. K. & Li, S. S. Non-histone protein methylation as a regulator of cellular signalling and function. Nat Rev Mol Cell Biol 16, 5–17 (2015). 10.1038/nrm3915

8 Guo, J. et al. AKT methylation by SETDB1 promotes AKT kinase activity and oncogenic functions. Nat Cell Biol 21, 226–237 (2019). 10.1038/s41556-018-0261-6

9 Wang, G. et al. SETDB1-mediated methylation of Akt promotes its K63-linked ubiquitination and activation leading to tumorigenesis. Nat Cell Biol 21, 214–225 (2019). 10.1038/s41556-018-0266-1

10 Chan, L. H. et al. PRMT6 Regulates RAS/RAF Binding and MEK/ERK-Mediated Cancer Stemness Activities in Hepatocellular Carcinoma through CRAF Methylation. Cell Rep 25, 690–701 e698 (2018). 10.1016/j.celrep.2018.09.053

11 Kim, E. et al. Phosphorylation of EZH2 activates STAT3 signaling via STAT3 methylation and promotes tumorigenicity of glioblastoma stem-like cells. Cancer Cell 23, 839–852 (2013). 10.1016/j.ccr.2013.04.008

12 He, A. et al. PRC2 directly methylates GATA4 and represses its transcriptional activity. Genes Dev 26, 37–42 (2012). 10.1101/gad.173930.111

13 Park, S. H. et al. Posttranslational regulation of FOXA1 by Polycomb and BUB3/USP7 deubiquitin complex in prostate cancer. Sci Adv 7 (2021). 10.1126/sciadv.abe2261

14 Hoy, S. M. Tazemetostat: First Approval. Drugs 80, 513–521 (2020). 10.1007/s40265-020-01288-x

15 Siu LL R. D., Vinay SP, Romano PM, Menis J, Opdam FL, Heinhuis KM, Egger JL, Gorman SA, Parasrampuria R, Wang K, Kremer BE, Gounder MM. METEOR-1: A phase I study of GSK3326595, a first-in-class protein arginine methyltransferase 5 (PRMT5) inhibitor, in advanced solid tumours. Annals of Oncology 30 (2019).

16 Watts JM B. T., Thomassen A, Brunner AW, Minden MD, Papadantonakis N, Abedin S, Baines AJ, Barbash O, Gorman S, Kremer BE, Borthakur GM. A Phase I/II Study to Investigate the Safety and Clinical Activity of the Protein Arginine Methyltransferase 5 Inhibitor GSK3326595 in Subjects with Myelodysplastic Syndrome and Acute Myeloid Leukemia. Blood 134 (2019).

17 Ong, S. E., Mittler, G. & Mann, M. Identifying and quantifying in vivo methylation sites by heavy methyl SILAC. Nat Methods 1, 119–126 (2004). 10.1038/nmeth715

18 Cao, X. J., Arnaudo, A. M. & Garcia, B. A. Large-scale global identification of protein lysine methylation in vivo. Epigenetics 8, 477–485 (2013). 10.4161/epi.24547

19 Guo, A. et al. Immunoaffinity enrichment and mass spectrometry analysis of protein methylation. Mol Cell Proteomics 13, 372–387 (2014). 10.1074/mcp.O113.027870

20 Levy, D. et al. A proteomic approach for the identification of novel lysine methyltransferase substrates. Epigenetics Chromatin 4, 19 (2011). 10.1186/1756-8935-4-19

21 Bremang, M. et al. Mass spectrometry-based identification and characterisation of lysine and arginine methylation in the human proteome. Mol Biosyst 9, 2231–2247 (2013). 10.1039/c3mb00009e

22 Moore, K. E. & Gozani, O. An unexpected journey: lysine methylation across the proteome. Biochim Biophys Acta 1839, 1395–1403 (2014). 10.1016/j.bbagrm.2014.02.008

23 Hart-Smith, G., Yagoub, D., Tay, A. P., Pickford, R. & Wilkins, M. R. Large Scale Mass Spectrometry-based Identifications of Enzyme-mediated Protein Methylation Are Subject to High False Discovery Rates. Mol Cell Proteomics 15, 989–1006 (2016). 10.1074/mcp.M115.055384

24 Nwajiobi, O., Mahesh, S., Streety, X. & Raj, M. Selective Triazenation Reaction (STaR) of Secondary Amines for Tagging Monomethyl Lysine Post-Translational Modifications. Angew Chem Int Ed Engl 60, 7344–7352 (2021). 10.1002/anie.202013997

25 Chalker, J. M., Bernardes, G. J. & Davis, B. G. A “tag-and-modify” approach to site-selective protein modification. Acc Chem Res 44, 730–741 (2011). 10.1021/ar200056q

26 Islam, K. The Bump-and-Hole Tactic: Expanding the Scope of Chemical Genetics. Cell Chem Biol 25, 1171–1184 (2018). 10.1016/j.chembiol.2018.07.001

27 Wang, R. et al. Profiling genome-wide chromatin methylation with engineered posttranslation apparatus within living cells. J Am Chem Soc 135, 1048–1056 (2013). 10.1021/ja309412s

28 Wang, R. & Luo, M. A journey toward Bioorthogonal Profiling of Protein Methylation inside living cells. Curr Opin Chem Biol 17, 729–737 (2013). 10.1016/j.cbpa.2013.08.007

29 Peters, W. et al. Enzymatic site-specific functionalization of protein methyltransferase substrates with alkynes for click labeling. Angew Chem Int Ed Engl 49, 5170–5173 (2010). 10.1002/anie.201001240

30 Bothwell, I. R. et al. Se-adenosyl-L-selenomethionine cofactor analogue as a reporter of protein methylation. J Am Chem Soc 134, 14905–14912 (2012). 10.1021/ja304782r

31 Sohtome, Y., Shimazu, T., Shinkai, Y. & Sodeoka, M. Propargylic Se-adenosyl-l-selenomethionine: A Chemical Tool for Methylome Analysis. Acc Chem Res 54, 3818–3827 (2021). 10.1021/acs.accounts.1c00395

32 Davydova, E. et al. The methyltransferase METTL9 mediates pervasive 1-methylhistidine modification in mammalian proteomes. Nat Commun 12, 891 (2021). 10.1038/s41467-020-20670-7

33 Hartstock, K. et al. Enzymatic or In Vivo Installation of Propargyl Groups in Combination with Click Chemistry for the Enrichment and Detection of Methyltransferase Target Sites in RNA. Angew Chem Int Ed Engl 57, 6342–6346 (2018). 10.1002/anie.201800188

34 Muttach, F. & Rentmeister, A. A Biocatalytic Cascade for Versatile One-Pot Modification of mRNA Starting from Methionine Analogues. Angew Chem Int Ed Engl 55, 1917–1920 (2016). 10.1002/anie.201507577

35 Mikutis, S. et al. meCLICK-Seq, a Substrate-Hijacking and RNA Degradation Strategy for the Study of RNA Methylation. ACS Cent Sci 6, 2196–2208 (2020). 10.1021/acscentsci.0c01094

36 Knutson, S. K. et al. Durable tumor regression in genetically altered malignant rhabdoid tumors by inhibition of methyltransferase EZH2. Proc Natl Acad Sci U S A 110, 7922–7927 (2013). 10.1073/pnas.1303800110

37 Dieterich, D. C., Link, A. J., Graumann, J., Tirrell, D. A. & Schuman, E. M. Selective identification of newly synthesized proteins in mammalian cells using bioorthogonal noncanonical amino acid tagging (BONCAT). Proc Natl Acad Sci U S A 103, 9482–9487 (2006). 10.1073/pnas.0601637103

38 Hornbeck, P. V. et al. PhosphoSitePlus, 2014: mutations, PTMs and recalibrations. Nucleic Acids Res 43, D512–520 (2015). 10.1093/nar/gku1267

39 Krassowski, M. et al. ActiveDriverDB: human disease mutations and genome variation in post-translational modification sites of proteins. Nucleic Acids Res 46, D901–D910 (2018). 10.1093/nar/gkx973

40 Menendez, D., Inga, A. & Resnick, M. A. The expanding universe of p53 targets. Nat Rev Cancer 9, 724–737 (2009). 10.1038/nrc2730

41 Jansson, M. et al. Arginine methylation regulates the p53 response. Nat Cell Biol 10, 1431–1439 (2008). 10.1038/ncb1802

42 Chuikov, S. et al. Regulation of p53 activity through lysine methylation. Nature 432, 353–360 (2004). 10.1038/nature03117

43 Huang, J. et al. Repression of p53 activity by Smyd2-mediated methylation. Nature 444, 629–632 (2006). 10.1038/nature05287

44 Fong, J. Y. et al. Therapeutic Targeting of RNA Splicing Catalysis through Inhibition of Protein Arginine Methylation. Cancer Cell 36, 194–209 e199 (2019). 10.1016/j.ccell.2019.07.003

45 Meister, G. et al. Methylation of Sm proteins by a complex containing PRMT5 and the putative U snRNP assembly factor pICln. Curr Biol 11, 1990–1994 (2001). 10.1016/s0960-9822(01)00592-9

46 Bezzi, M. et al. Regulation of constitutive and alternative splicing by PRMT5 reveals a role for Mdm4 pre-mRNA in sensing defects in the spliceosomal machinery. Genes Dev 27, 1903–1916 (2013). 10.1101/gad.219899.113

47 Hodge, R. G. & Ridley, A. J. Regulating Rho GTPases and their regulators. Nat Rev Mol Cell Biol 17, 496–510 (2016). 10.1038/nrm.2016.67

48 Bikkavilli, R. K. & Malbon, C. C. Wnt3a-stimulated LRP6 phosphorylation is dependent upon arginine methylation of G3BP2. J Cell Sci 125, 2446–2456 (2012). 10.1242/jcs.100933

49 Stricher, F., Macri, C., Ruff, M. & Muller, S. HSPA8/HSC70 chaperone protein: structure, function, and chemical targeting. Autophagy 9, 1937–1954 (2013). 10.4161/auto.26448

50 Szewczyk, M. M. et al. Pharmacological inhibition of PRMT7 links arginine monomethylation to the cellular stress response. Nat Commun 11, 2396 (2020). 10.1038/s41467-020-16271-z

51 Gao, W. W. et al. Arginine methylation of HSP70 regulates retinoid acid-mediated RARbeta2 gene activation. Proc Natl Acad Sci U S A 112, E3327–3336 (2015). 10.1073/pnas.1509658112

52 Brown-Borg, H. M. Reduced growth hormone signaling and methionine restriction: interventions that improve metabolic health and extend life span. Ann N Y Acad Sci 1363, 40–49 (2016). 10.1111/nyas.12971

53 Hu, C. et al. Heat shock proteins: Biological functions, pathological roles, and therapeutic opportunities. MedComm (2020) 3, e161 (2022). 10.1002/mco2.161

54 Deng, W. et al. Arginine methylation of HSPA8 by PRMT9 inhibits ferroptosis to accelerate hepatitis B virus-associated hepatocellular carcinoma progression. J Transl Med 21, 625 (2023). 10.1186/s12967-023-04408-9

55 Yin, X. K. et al. PRMT1 enhances oncogenic arginine methylation of NONO in colorectal cancer. Oncogene 40, 1375–1389 (2021). 10.1038/s41388-020-01617-0

56 A, P., et al. EZH2 promotes DNA replication by stabilizing interaction of POLdelta and PCNA via methylation-mediated PCNA trimerization. Epigenetics Chromatin 11, 44 (2018). 10.1186/s13072-018-0213-1

57 Hwang, J., Lee, J. A. & Pallas, D. C. Leucine Carboxyl Methyltransferase 1 (LCMT-1) Methylates Protein Phosphatase 4 (PP4) and Protein Phosphatase 6 (PP6) and Differentially Regulates the Stable Formation of Different PP4 Holoenzymes. J Biol Chem 291, 21008–21019 (2016). 10.1074/jbc.M116.739920

58 Singh, S. et al. Facile chemoenzymatic strategies for the synthesis and utilization of S-adenosyl-(L)-methionine analogues. Angew Chem Int Ed Engl 53, 3965–3969 (2014). 10.1002/anie.201308272

59 Massignani, E. et al. ProMetheusDB: An In-Depth Analysis of the High-Quality Human Methyl-proteome. Mol Cell Proteomics 21, 100243 (2022). 10.1016/j.mcpro.2022.100243

60 Hartstock, K. et al. Enzymatic or in vivo installation of propargyl groups in combination with click chemistry for the enrichment and detection of methyltransferase target sites in RNA. Angewandte Chemie International Edition 57, 6342--6346 (2018).

61 Zhou, Y. et al. Metascape provides a biologist-oriented resource for the analysis of systems-level datasets. Nat Commun 10, 1523 (2019). 10.1038/s41467-019-09234-6

62 Soucek, S. et al. Evolutionarily conserved polyadenosine RNA binding protein Nab2 cooperates with splicing machinery to regulate the fate of pre-mRNA. Molecular and cellular biology 36, 2697–2714 (2016).

63 Seyfried, N. T. et al. A multi-network approach identifies protein-specific co-expression in asymptomatic and symptomatic Alzheimer’s disease. Cell systems 4, 60–72. e64 (2017).

64 Zhu, Y. et al. DEqMS: a method for accurate variance estimation in differential protein expression analysis. Molecular & Cellular Proteomics 19, 1047–1057 (2020).

65 Hornbeck, P. V. et al. PhosphoSitePlus, 2014: mutations, PTMs and recalibrations. Nucleic acids research 43, D512–D520 (2015).

66 Krassowski, M. et al. ActiveDriverDB: human disease mutations and genome variation in post-translational modification sites of proteins. Nucleic acids research 46, D901–D910 (2018).

