## Supplementary material for "Dynamic *in vivo* mapping of the methylproteome using a chemoenzymatic approach": PSM Sup Info 5_12_24.pdf

**Extended Data**

Supplementary Tables

**Tables**

**Supplementary Table 1**: CuAAC reaction conditions for experimental applications.

| Application | $CuSO_4$ | THPTA | NaAsc | $N_3$ | DMSO | TX-100 | Time |
|---|---|---|---|---|---|---|---|
| WB | 1 mM | 1.5 mM | 3 mM | 25 µM | None | None | 30 m |
| IF | 1 mM | 1.5 mM | 3 mM | 25 µM | None | None | 30 m |
| IHC | 1 mM | 1.5 mM | 3 mM | 25 µM | 5% | 0.2% | 45 m |
| Biotin IP | 1 mM | 1.5 mM | 3 mM | 200 µM | None | None | 1 h |

**Extended Figures**

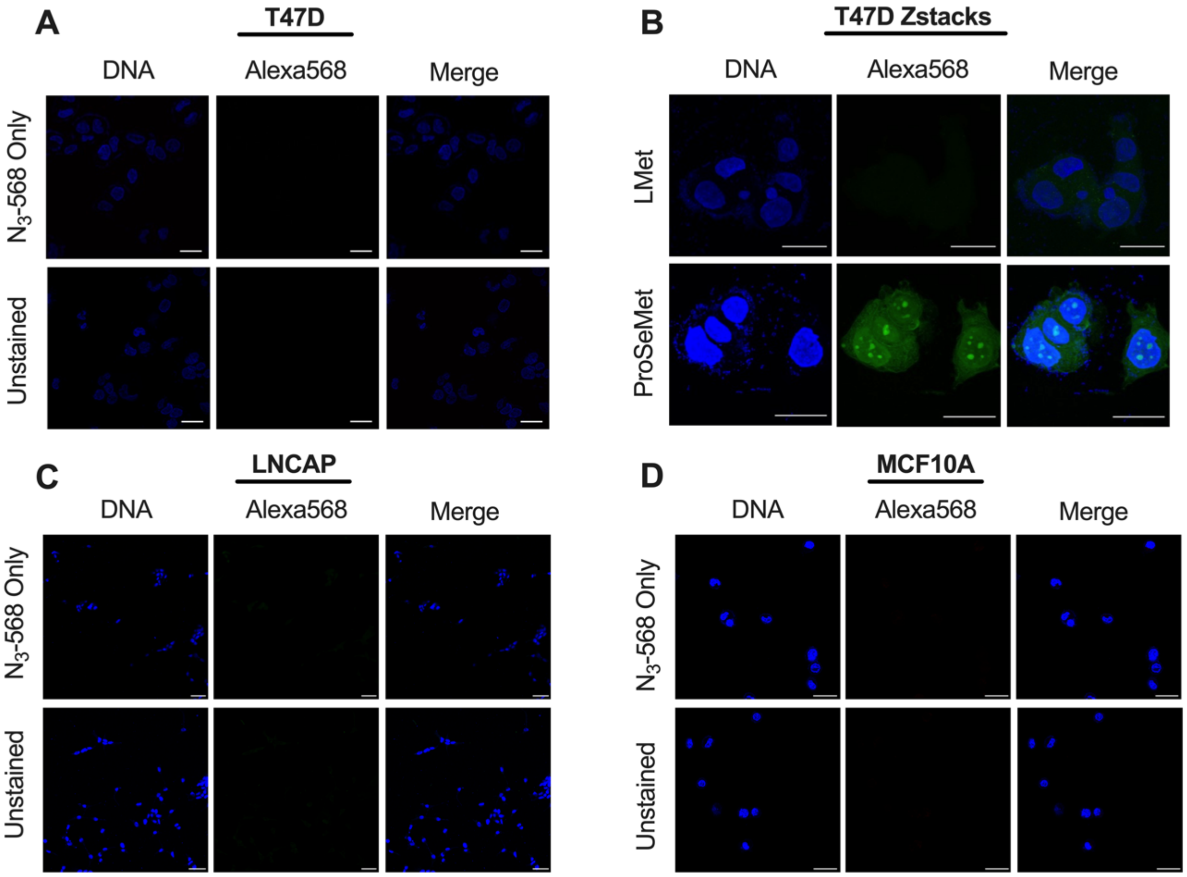


**Supplementary Figure 1. IF controls for ProSeMet/L-Met labeling. IF controls for ProSeMet/L-Met labeling.** T47D (**a**), LNCaP (**c**), and MCF10A (**d**) cells were treated with 100 µM ProSeMet or L-Met 16 h, fixed, permeabilized, and incubated with fluorescent picolyl azide alone or Hoechst stain alone. No nonspecific binding is observed in T47D, LNCaP, and MCF10A. Scale bar represents 25 µm (T47D, MCF10A) or 50 µM (LNCaP). **b.** T47D cells treated with ProSeMet or L-Met 16 h were fixed, permeabilized, and subjected to CuAAC to attach a fluorescent picolyl azide. Zstack image represents maximal intensity of each slice.

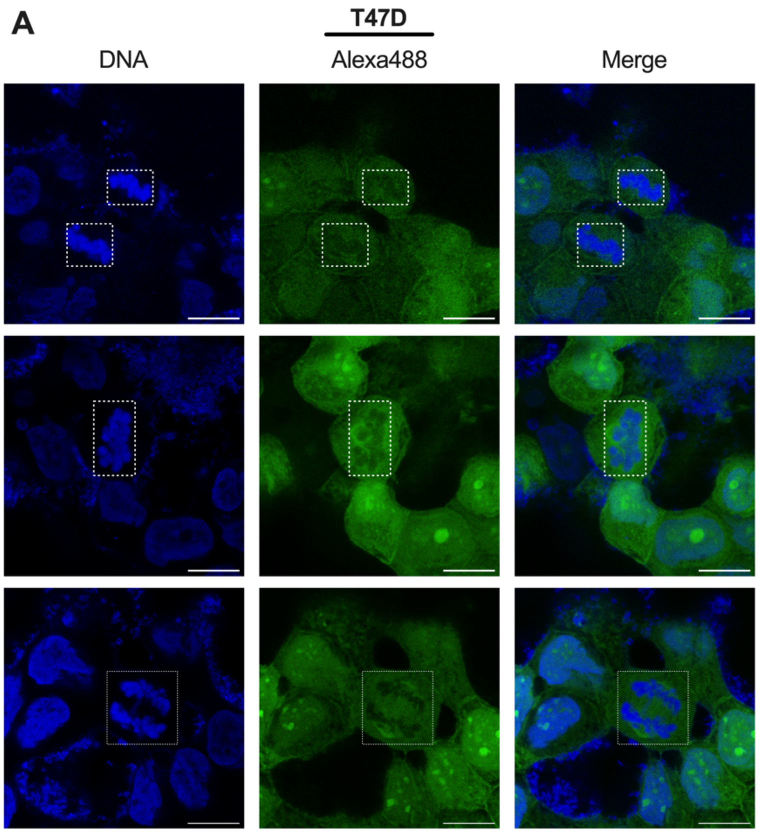


**Supplementary Figure 2. ProSeMet does not label condensed DNA.** T47D cells treated with 100 μM ProSeMet 16 h were fixed, permeabilized, and subjected to CuAAC to attach a fluorescent picolyl azide. **a**. Condensed DNA observed during cellular division (top: cytokinesis, middle: prophase, bottom: anaphase).

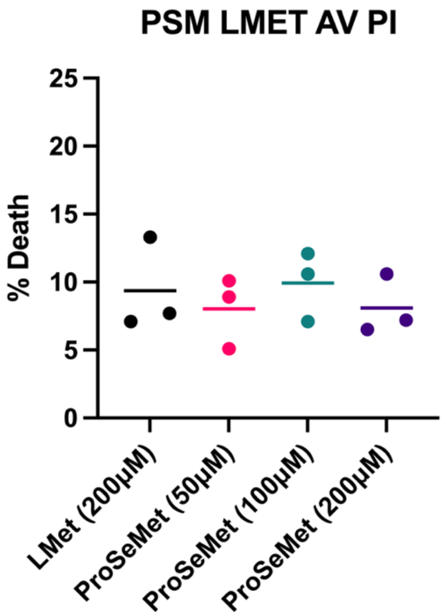


**Supplementary Figure 3. ProSeMet treatment does not cause cell death.** T47D cells treated with 50-200 μM ProSeMet for 24 h were stained with Annexin V/PI and analyzed by flow cytometry (*n=3*).

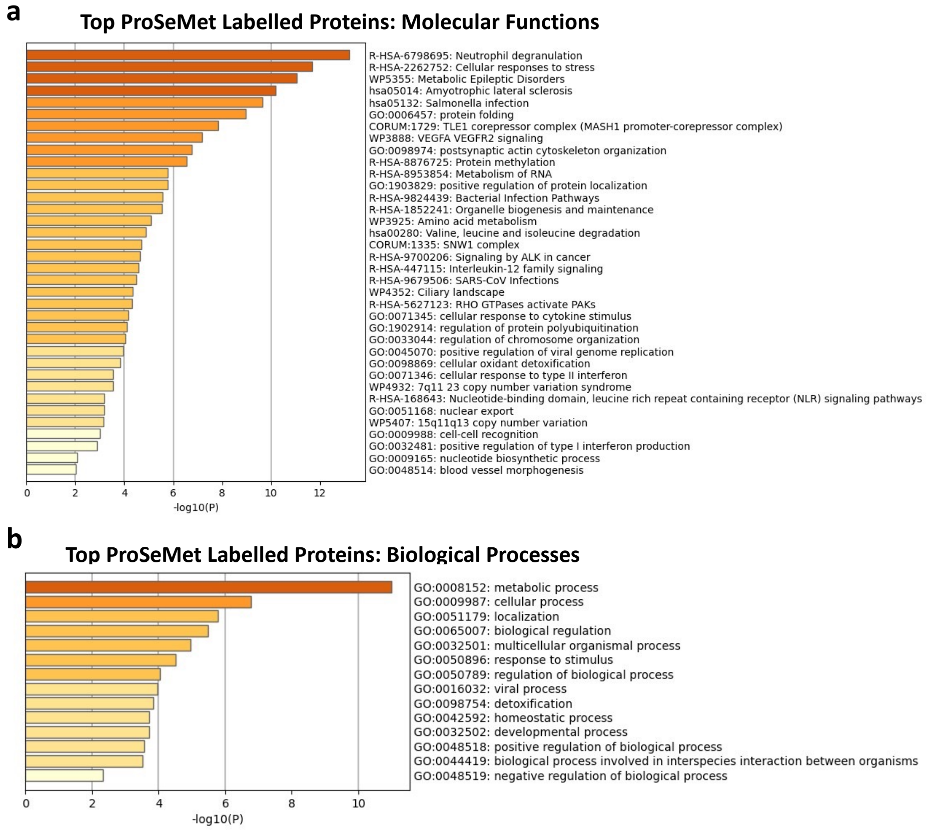


**Supplementary Figure 4. Gene Ontology (GO) analysis of ProSeMet sites from unenriched LC-MS/MS in mammalian cells.** (**a**) and (**b**). A gene list of propargylated proteins was utilized as input in metascape, with input and analysis species set to H. sapiens. Pathway and process enrichment analysis was carried out with the following ontology sources: KEGG Pathway, GO Biological Processes, Reactome Gene Sets, Canonical Pathways, CORUM, WikiPathways, and PANTHER Pathway. All genes in the human genome were used as the enrichment background. Terms with a p-value $< 0.01$, a minimum count of 3, and an enrichment factor $> 1.5$ were utilized. p-values were calculated based on the cumulative hypergeometric distribution, and q-values are calculated using the Benjamini-Hochberg procedure. The best term within a group is chosen as the group summary. ***Excel sheet containing analysis can be found in Supplementary data 2.***

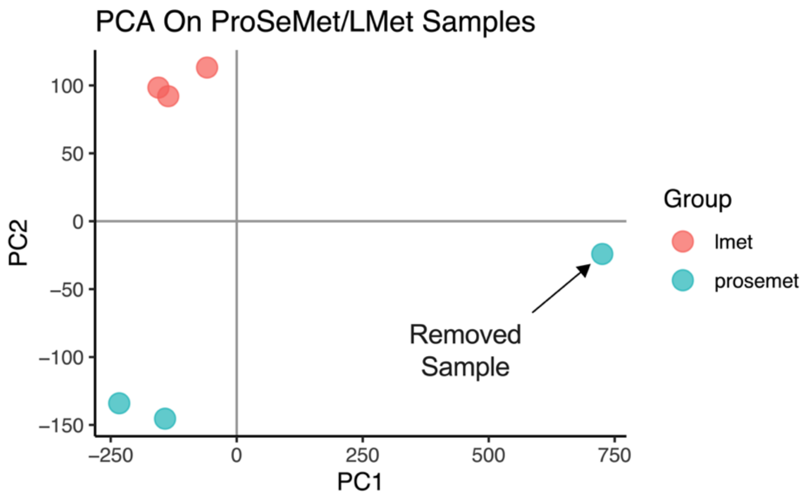


**Supplementary Figure 5. PCA analysis of biological replicates submitted for LC-MS/MS analysis.** PCA analysis of biological replicates submitted for LC-MS/MS analysis, generated via statistical software. ProSeMet sample 3 was removed prior to further analysis.

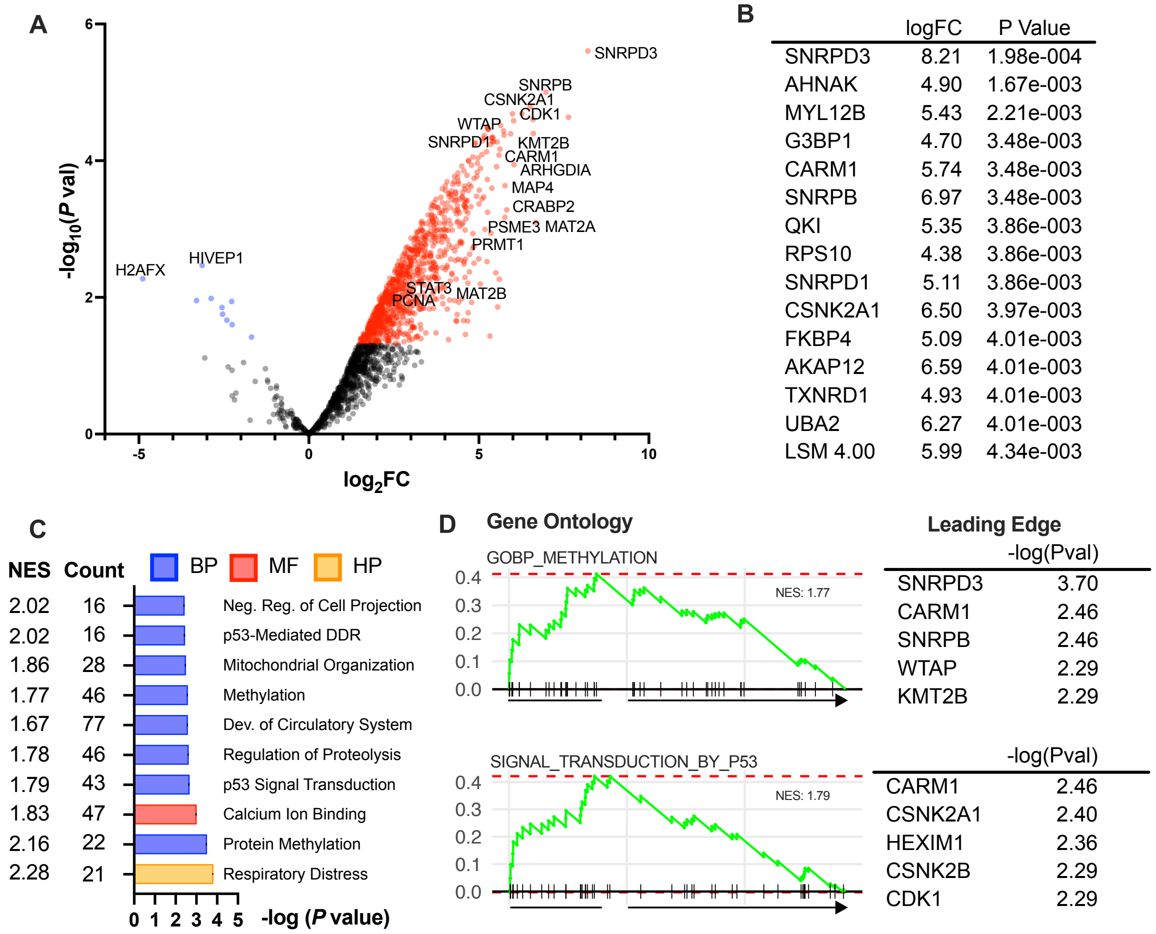


**Supplementary Figure 6. Pseudomethylated proteins converge into predicted gene ontologies while extending the methylproteome.** G401 cells treated with 100 μM ProSeMet or 100 μM L-Met 16 h were lysed, subjected to a click reaction to attach a biotin-conjugated picolyl azide, and enriched using magnetic streptavidin beads. Enriched proteins were digested with trypsin and analyzed via label-free LC-MS/MS. **a.** Volcano plot showing statistical significance vs $\log_2$-fold change (logFC) of proteins isolated from G401 cells treated with ProSeMet or L-Met. Data represents 3 biological replicates consisting of matched experiments. Statistical significance was calculated using a two-tailed unpaired *t*-test. **b**. Top 15 enriched proteins. Statistical significance represents scaled *P* values after differential expression analysis. **c**. Gene set enrichment analysis (GSEA) of ProSeMet enriched proteins using GO gene set. The top ten pathways are shown with corresponding number of proteins (count) and normalized enrichment score (NES). **d**. GSEA analysis of methylation (top) and p53-mediated signal transduction (bottom) gene sets. Top 5 enriched proteins by *P* value in the leading edge are shown to the right. For **c** and **d**, statistical significance was assessed using Fast Gene Set Enrichment Analysis (fgsea).

A

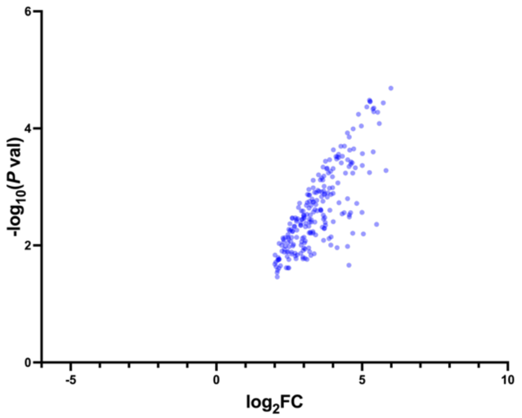


| | logFC | logPValue |
|---|---|---|
| LSM 4.00 | 5.989810 | 4.686916 |
| S100A1 | 5.256397 | 4.482155 |
| CDK1 | 5.280067 | 4.460858 |
| EPHA2 | 5.275312 | 4.454965 |
| HMOX2 | 5.728186 | 4.437184 |
| WTAP | 5.163824 | 4.366803 |
| IFITM3 | 5.406214 | 4.348426 |
| ISG15 | 5.373568 | 4.331429 |
| CARS | 5.399502 | 4.284220 |
| TNFRSF10B | 5.533544 | 4.275322 |

B

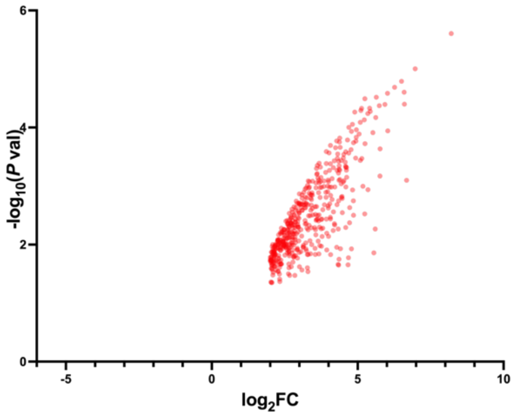


| | logFC | logPValue |
|---|---|---|
| SNRPD3 | 8.206059 | 5.606909 |
| SNRPB | 6.973263 | 5.004295 |
| CSNK2A1 | 6.502378 | 4.789590 |
| UBA2 | 6.265811 | 4.686400 |
| AKAP12 | 6.594047 | 4.603001 |
| PABPN1 | 6.021634 | 4.585162 |
| RPA2 | 5.640058 | 4.517992 |
| HIGD2A | 5.245443 | 4.492654 |
| HEXIM1 | 6.604102 | 4.399010 |
| KMT2B | 5.933474 | 4.393224 |

**Supplementary Figure 7. Analysis of ProSeMet-labeled proteome identified 707 proteins with uncharacterized methylation sites using PhosphoSitePlus database.** Significantly enriched proteins identified *via* LC-MS/MS (p-value < 0.05, logFC > 2) were screened against the PhosphoSitePlus database of proteins with known methylation sites to identify (**a**) 221 proteins with uncharacterized methylation sites and (**b**) 486 proteins with previously identified methylation sites. Table to the right of each volcano plot indicates the top ten identified genes by p value.

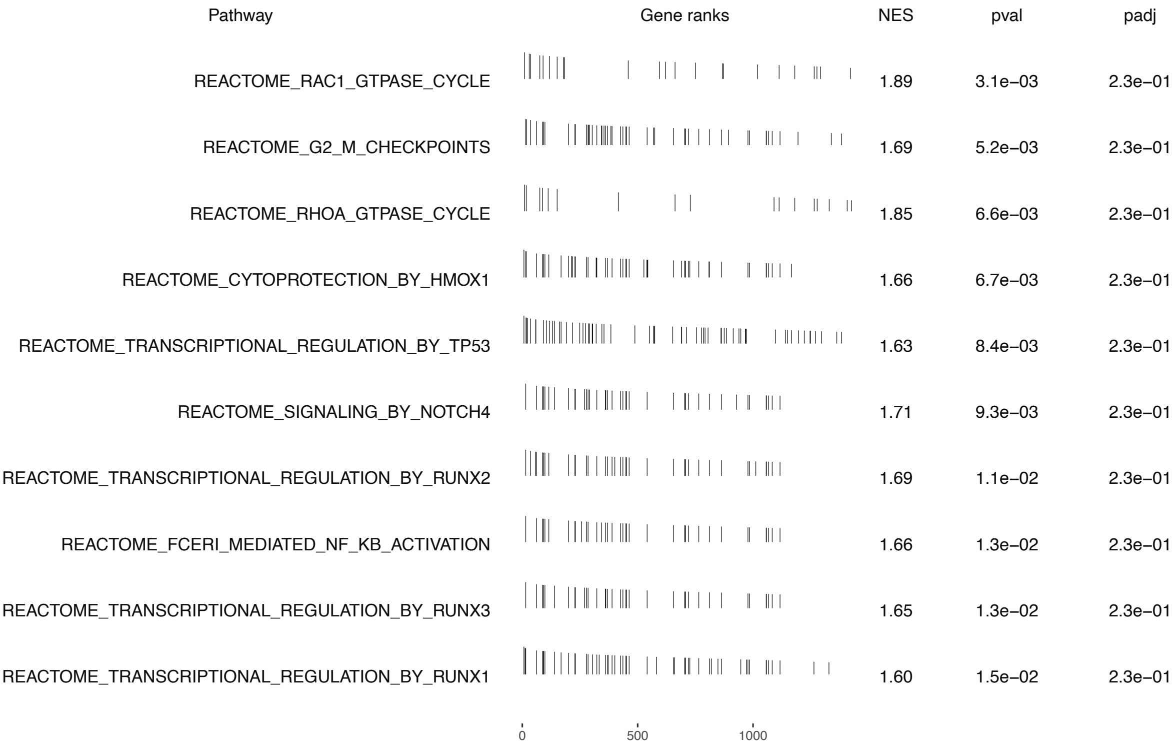


**Supplementary Figure 8. GSEA analysis of ProSeMet enriched proteins using REACTOME molecular signatures database.** GSEA analysis of ProSeMet enriched proteins using REACTOME gene set. The top ten pathways are shown with gene distribution based on rankings, normalized enrichment score (NES), *P* value, and adjusted *P* value (calculated via statistical software).

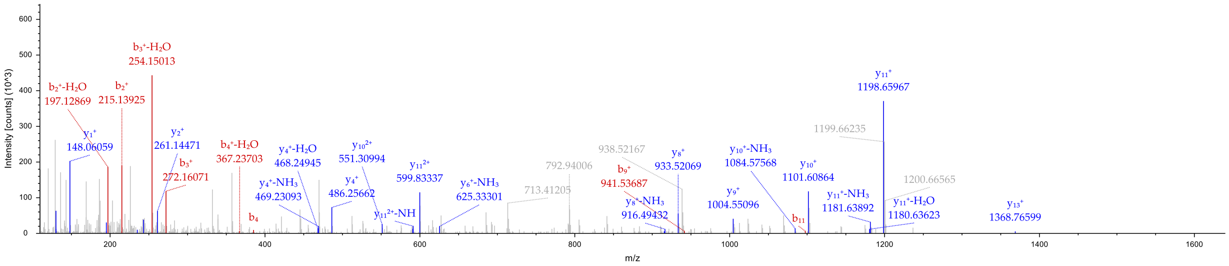


| #1 | b+ | b2+ | Seq. | y+ | y2+ | #2 |
|---|---|---|---|---|---|---|
| 1 | 114.09134 | 57.54931 | L | | | 15 |
| 2 | 215.13902 | 108.07315 | T | 1469.811... | 735.40921 | 14 |
| 3 | 272.16048 | 136.58388 | G | 1368.763... | 684.88537 | 13 |
| 4 | 385.24455 | 193.12591 | I | 1311.742... | 656.37464 | 12 |
| 5 | 482.29731 | 241.65229 | P | 1198.657... | 599.83260 | 11 |
| 6 | 579.35007 | 290.17868 | P | 1101.605... | 551.30622 | 10 |
| 7 | 650.38719 | 325.69723 | A | 1004.552... | 502.77984 | 9 |
| 8 | 747.43995 | 374.22361 | P | 933.51529 | 467.26128 | 8 |
| 9 | 941.55676 | 471.28202 | R-Ben... | 836.46253 | 418.73490 | 7 |
| 10 | 998.57823 | 499.79275 | G | 642.34572 | 321.67650 | 6 |
| 11 | 1097.646... | 549.32696 | V | 585.32425 | 293.16576 | 5 |
| 12 | 1194.699... | 597.85334 | P | 486.25584 | 243.63156 | 4 |
| 13 | 1322.757... | 661.88263 | Q | 389.20308 | 195.10518 | 3 |
| 14 | 1435.842... | 718.42466 | I | 261.14450 | 131.07589 | 2 |
| 15 | | | E | 148.06043 | 74.53386 | 1 |

**Supplementary Figure 9. HSPA8 R469 is propargylated.** Representative peptide spectra of modified peptides R469 propargylation (propargyl arginine site, +38 on arginine). Sequence: LTGIPPAPRGVPQIE, R9-BenProsemet1 (38.01570 Da).

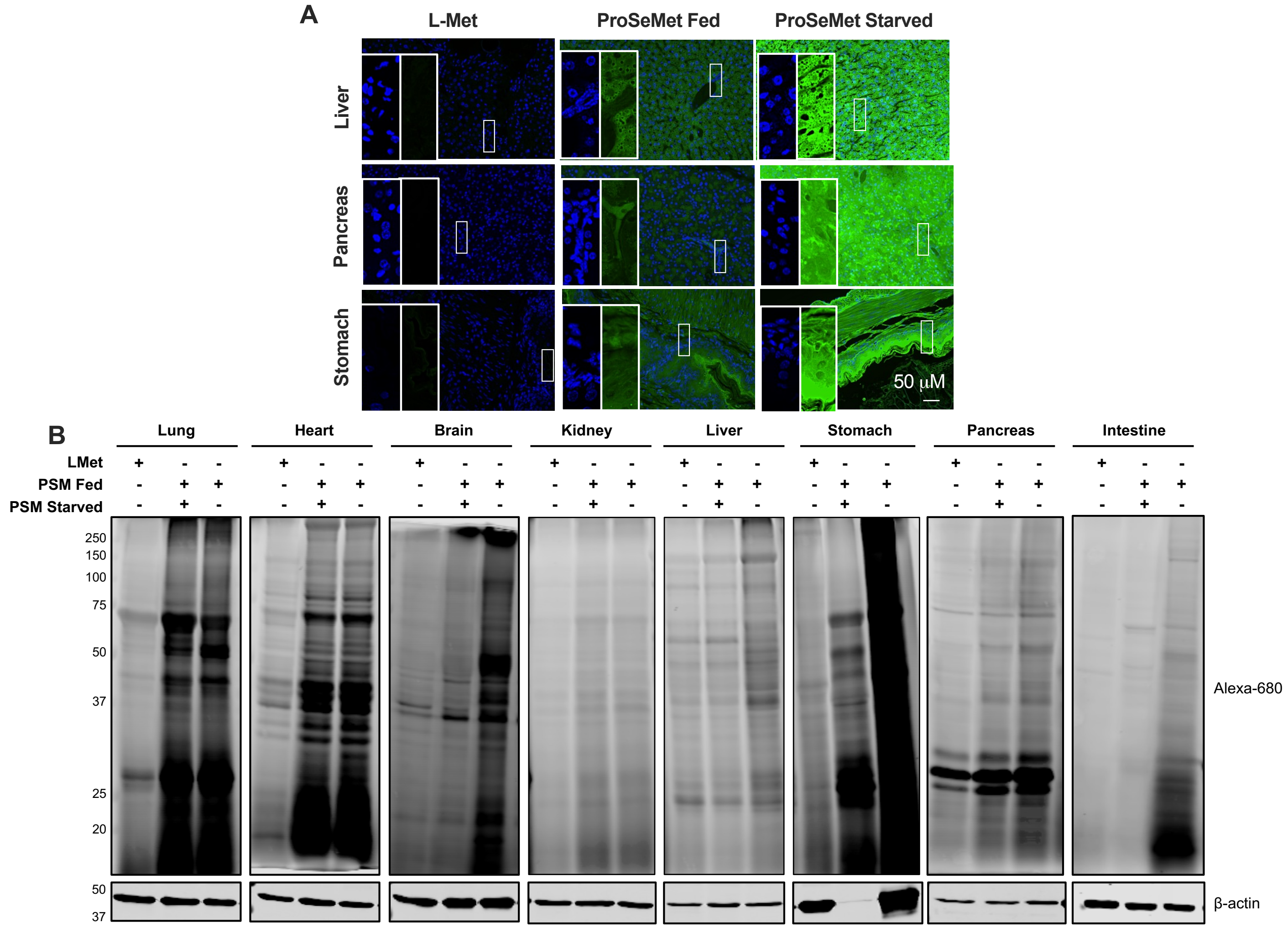


**Supplementary Figure 10. In-tissue fluorescence and blot analysis of organs collected from mouse treated with ProSeMet or L-Met.** Mice treated with ProSeMet or L-Met in the presence of food or starved were euthanized and organs were collected. **a.** Organs extracted from mice treated with 15 mg ProSeMet or equimolar L-Met via IP injection while being fed or starved for 12 h were formaldehyde fixed and paraffin embedded. Tissue sections were subjected to CuAAC to attach a fluorescent picolyl azide (568 nm, pseudocolored green) and counterstained with DAPI (blue). In-tissue fluorescence analysis of liver (top), pancreas (middle), and stomach (bottom), $n \geq 3$. **b.** Representative blot images to accompany the densitometry shown in Figure 5h, $n \geq 3$.

**a**

### Top ProSeMet Labelled Proteins: Brain

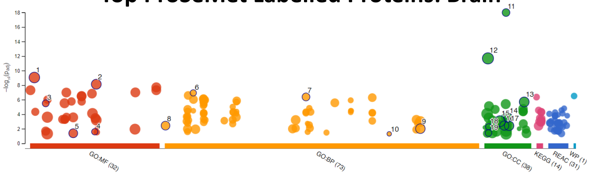


| ID | Source | Term ID | Term Name | $p_{adj}$ (query_1) |
|---|---|---|---|---|
| 1 | GO:MF | GO:0003824 | catalytic activity | $8.923\times10^{-10}$ |
| 2 | GO:MF | GO:0043168 | anion binding | $7.488\times10^{-9}$ |
| 3 | GO:MF | GO:0005200 | structural constituent of cytoskeleton | $3.202\times10^{-6}$ |
| 4 | GO:MF | GO:0042288 | MHC class I protein binding | $2.588\times10^{-2}$ |
| 5 | GO:MF | GO:0019901 | protein kinase binding | $4.160\times10^{-2}$ |
| 6 | GO:BP | GO:0006734 | NADH metabolic process | $1.163\times10^{-7}$ |
| 7 | GO:BP | GO:0046034 | ATP metabolic process | $4.167\times10^{-7}$ |
| 8 | GO:BP | GO:0000226 | microtubule cytoskeleton organization | $3.685\times10^{-3}$ |
| 9 | GO:BP | GO:1901566 | organonitrogen compound biosynthetic process | $9.718\times10^{-3}$ |
| 10 | GO:BP | GO:0098735 | positive regulation of the force of heart contraction | $4.995\times10^{-2}$ |
| 11 | GO:CC | GO:0043209 | myelin sheath | $1.036\times10^{-18}$ |
| 12 | GO:CC | GO:0005737 | cytoplasm | $2.081\times10^{-12}$ |
| 13 | GO:CC | GO:0120025 | plasma membrane bounded cell projection | $2.034\times10^{-6}$ |
| 14 | GO:CC | GO:0044297 | cell body | $2.916\times10^{-4}$ |
| 15 | GO:CC | GO:0032991 | protein-containing complex | $9.028\times10^{-4}$ |
| 16 | GO:CC | GO:0036477 | somatodendritic compartment | $3.625\times10^{-3}$ |
| 17 | GO:CC | GO:0045202 | synapse | $3.890\times10^{-3}$ |
| 18 | GO:CC | GO:0005790 | smooth endoplasmic reticulum | $5.285\times10^{-3}$ |
| 19 | GO:CC | GO:0005791 | rough endoplasmic reticulum | $3.885\times10^{-2}$ |

**b**

### Top ProSeMet Labelled Proteins: Heart

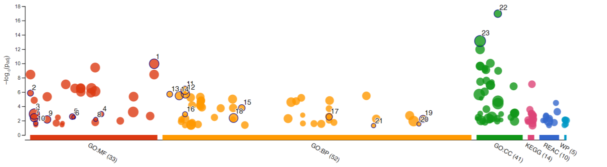


| ID | Source | Term ID | Term Name ↑ | $p_{adj}$ (query_1) |
|---|---|---|---|---|
| 6 | GO:MF | GO:0030060 | L-malate dehydrogenase activity | $3.340\times10^{-3}$ |
| 10 | GO:MF | GO:0003985 | acetyl-CoA C-acetyltransferase activity | $2.328\times10^{-2}$ |
| 7 | GO:MF | GO:0003779 | actin binding | $3.928\times10^{-3}$ |
| 8 | GO:MF | GO:0043532 | angiostatin binding | $6.672\times10^{-3}$ |
| 9 | GO:MF | GO:0005516 | calmodulin binding | $6.838\times10^{-3}$ |
| 18 | GO:BP | GO:0019752 | carboxylic acid metabolic process | $4.333\times10^{-3}$ |
| 3 | GO:MF | GO:0003824 | catalytic activity | $1.198\times10^{-3}$ |
| 23 | GO:CC | GO:0005737 | cytoplasm | $7.446\times10^{-14}$ |
| 11 | GO:BP | GO:0006091 | generation of precursor metabolites and energy | $5.646\times10^{-7}$ |
| 1 | GO:MF | GO:1901363 | heterocyclic compound binding | $1.119\times10^{-10}$ |
| 21 | GO:BP | GO:0090131 | mesenchyme migration | $4.996\times10^{-2}$ |
| 2 | GO:MF | GO:0000146 | microfilament motor activity | $1.416\times10^{-6}$ |
| 14 | GO:BP | GO:0003012 | muscle system process | $3.315\times10^{-6}$ |
| 22 | GO:CC | GO:0043209 | myelin sheath | $1.098\times10^{-17}$ |
| 15 | GO:BP | GO:0030239 | myofibril assembly | $1.706\times10^{-4}$ |
| 20 | GO:BP | GO:1902109 | negative regulation of mitochondrial membrane p... | $3.001\times10^{-2}$ |
| 16 | GO:BP | GO:0006107 | oxaloacetate metabolic process | $1.277\times10^{-3}$ |
| 5 | GO:MF | GO:0019825 | oxygen binding | $2.745\times10^{-3}$ |
| 19 | GO:BP | GO:1902600 | proton transmembrane transport | $5.828\times10^{-3}$ |
| 4 | GO:MF | GO:0046933 | proton-transporting ATP synthase activity, rotation... | $1.219\times10^{-3}$ |
| 12 | GO:BP | GO:0006163 | purine nucleotide metabolic process | $1.827\times10^{-6}$ |
| 13 | GO:BP | GO:0002026 | regulation of the force of heart contraction | $2.037\times10^{-6}$ |
| 17 | GO:BP | GO:0055010 | ventricular cardiac muscle tissue morphogenesis | $3.041\times10^{-3}$ |

**c**

### Top ProSeMet Labelled Proteins: Lung

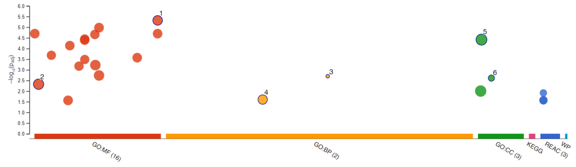


| ID | Source | Term ID | Term Name | $p_{adj}$ (query_1) |
|---|---|---|---|---|
| 1 | GO:MF | GO:1901363 | heterocyclic compound binding | $4.779\times10^{-6}$ |
| 2 | GO:MF | GO:0003824 | catalytic activity | $4.747\times10^{-3}$ |
| 3 | GO:BP | GO:0051659 | maintenance of mitochondrion location | $1.997\times10^{-3}$ |
| 4 | GO:BP | GO:0033554 | cellular response to stress | $2.495\times10^{-2}$ |
| 5 | GO:CC | GO:0005737 | cytoplasm | $3.834\times10^{-5}$ |
| 6 | GO:CC | GO:0031672 | A band | $2.449\times10^{-3}$ |

**Supplementary Figure 11. Gene Ontology (GO) analysis of ProSeMet sites from *in vivo* murine tissue.** Gene lists of propargylated proteins from murine brain (**a**), heart (**b**), and lung (**c**) tissue-derived LC-MS/MS were utilized as input in metascape and gProfiler, with input and analysis species set to *H. sapiens*. Pathway and process enrichment analysis was carried out with the following ontology sources: KEGG Pathway, GO Biological Processes, Reactome Gene Sets, Canonical Pathways, CORUM, WikiPathways, and PANTHER Pathway. All genes in the *M. Musculus* genome were used as the enrichment background. Terms with a p-value < 0.05, a minimum count of 3, and an enrichment factor > 1.5 were utilized. p-values were calculated based on the cumulative hypergeometric distribution, and q-values are calculated using the Benjamini-Hochberg procedure. ***Excel sheet containing analysis can be found in Supplementary data 4.***

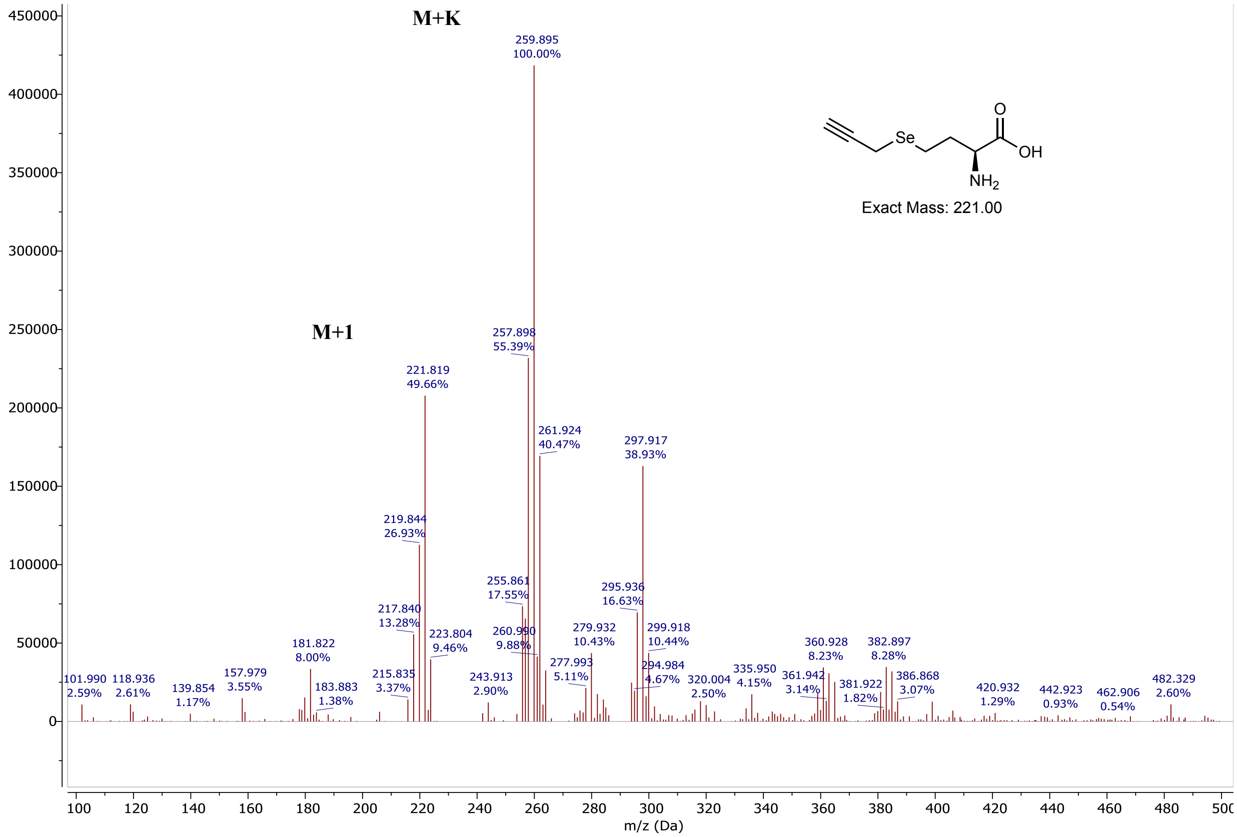


**Supplementary Figure 12. ESI-MS of synthesized ProSeMet.** Expected mass for $C_7H_{11}NO_2SeK^+$: 259.24 $[M + K]^+$; found: 259.815.

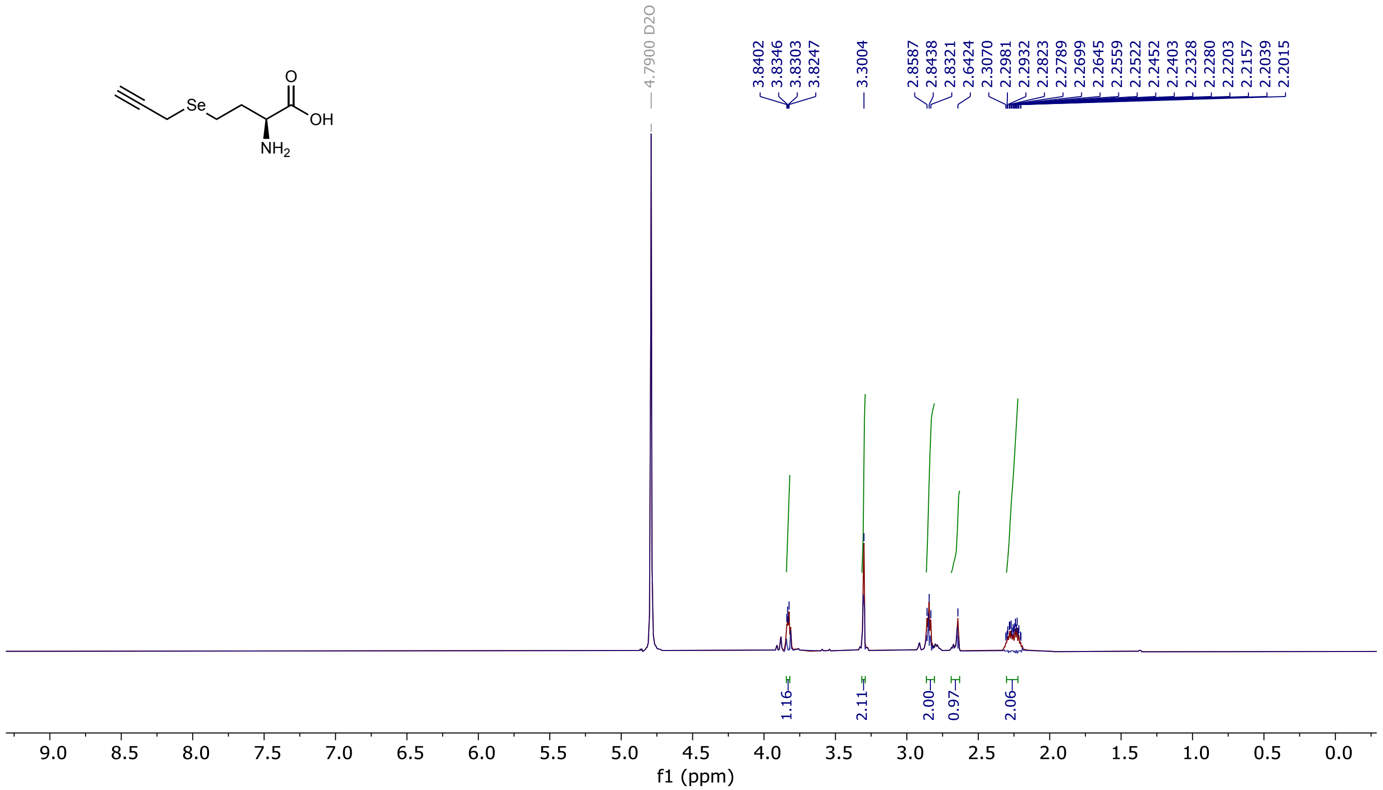


**Supplementary Figure 13. $^{1}H$ NMR of synthesized ProSeMet.** Purified ProSeMet was dissolved in dH2O and analyzed by $^{1}H$ NMR (600 MHz). $^{1}H$ NMR (600 MHz, D2O) δ 3.91-3.66 (m, 1H), 3.15 (s, 2H), 2.69 (dd, J = 9.1, 6.8 Hz, 2H), 2.49 (s, 1H), 2.14 – 2.07 (m, 2H).

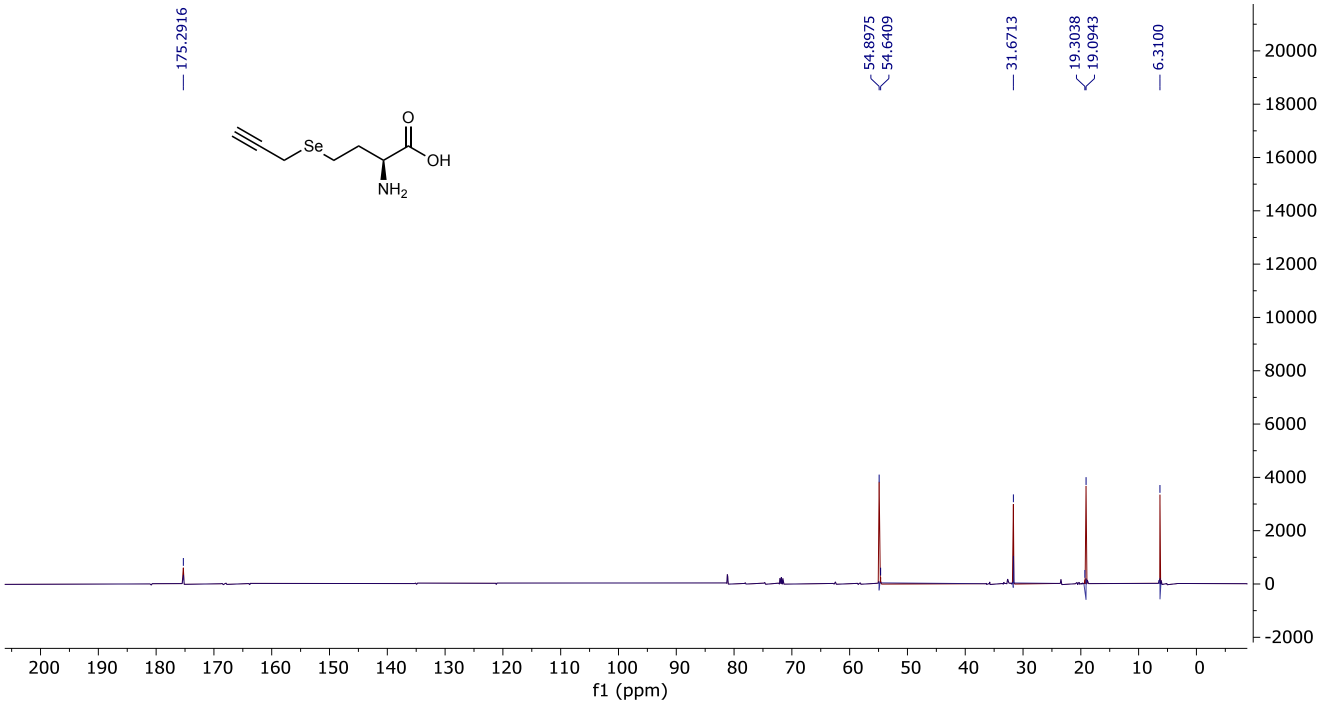


**Supplementary Figure 14. $^{13}C$ NMR of synthesized ProSeMet.** Purified ProSeMet was dissolved in dH2O and analyzed by $^{13}C$ NMR (151 MHz). $^{13}C$ NMR (151 MHz, D2O) δ 175.29, 54.90, 54.64, 31.67, 19.30, 19.09, 6.31.
